# Time-restricted feeding prevents memory impairments induced by obesogenic diet consumption in mice, in part through hippocampal thyroid hormone signaling

**DOI:** 10.1101/2024.04.29.591510

**Authors:** Jean-Christophe Helbling, Rachel Ginieis, Pierre Mortessagne, Mariano Ruiz-Gayo, Eva Gunnel Ducourneau, Dominique Ciocca, Illona-Marie Boulete, Alexandre Favereaux, Aurélia Ces, Enrica Montalban, Lucile Capuron, Freddy Jeanneteau, Guillaume Ferreira, Etienne Challet, Marie-Pierre Moisan

## Abstract

The consumption of calorie-rich diet has adverse effects on short and long-term memory, especially when introduced early in life when the brain is still maturing. Time-restricted feeding (TRF) without calorie restriction has proven to be an efficient strategy to reduce the deleterious effects of diet-induced obesity on metabolism. TRF was also shown to be beneficial to restore long-term memory in Alzheimer rodent models. Here, we provide evidence that four weeks of TRF restore the rhythmicity of some metabolic parameters together with short and long-term memory in mice fed a high fat-high sucrose (HFS) diet since weaning. Hippocampal translatome analyses indicated that impaired memory of mice under *ad libitum* HFS diet is accompanied by changes in genes associated with thyroid hormone signaling and astrocytic genes involved in the regulation of glutamate neurotransmission. TRF restored the diurnal expression variation of part of these genes and intra-hippocampal infusion of T3, the active form of thyroid hormone, rescued the memory performances of *ad libitum* HFS diet-fed mice. Thus, TRF demonstrates positive effects on both metabolism and memory in mice fed an obesogenic diet, highlighting this nutritional approach as a powerful tool in addressing obesity and its related comorbidities in mice. The analogous time-restricted eating in humans is an easy to implement lifestyle intervention that should now be tested in obese adolescents with memory alterations.

## Introduction

The prevalence of pediatric obesity has plateaued in most high-income countries but remains high, and it continues to rise in low and middle-income countries. Child and adolescent obesity are associated with multiple immediate and long-term negative health outcomes, impacting quality of life and increasing economic costs [1]. The consumption of an obesogenic diet, rich in saturated fat and refined sugar (HFS), at any time of day and night is one of the major causes of obesity, especially among adolescents [2]. HFS diet has deleterious consequences not only on cardiometabolic health but also on brain health and cognitive functions as shown by epidemiological and brain morphological analyses [3,4]. Over the last 15 years, data have accumulated in rodent models about the impact of consuming such diet on cognitive processes [5,6]. Early life consumption of HFS diet is particularly harmful for the brain that is still under development[7]. The hippocampus has been very much studied for its vulnerability to calorie-rich diet and its consequences on learning and memory. Our group has developed a model of juvenile HFS consumption in rodents, from weaning to adulthood and showed that this paradigm induced memory deficits [8,9]. Importantly, we found that HFS diet impaired hippocampal-dependent memories when started in juveniles, but did not affect memory when started at adulthood [8], through aberrant activity of the hippocampus [10,11]. Specifically, such memory impairments have been found to be associated with higher c-fos expression in the hippocampus and hippocampal efferent pathways to striatal and prefrontal areas [10,11] as well as aberrant synaptic transmission and plasticity in CA1 [12]. Interestingly, chemogenetic manipulation of the hippocampus or its efferent pathways rescued memory deficits for object location and long-term object recognition [11,13]. However, the molecular mechanism underpinnings of the HFS diet-induced memory impairments have not been identified yet.

HFS diet is also known to disturb circadian rhythms at the behavioral and molecular levels, favoring metabolic disturbances [14–17]. In particular, unlimited access to HFS food leads to an increased food intake during the resting phase and a reprogramming of the circadian expression of the transcriptome and metabolome in mice liver [18], suprachiasmatic nucleus of the hypothalamus and in prefrontal cortex [19]. The impact of juvenile HFS diet consumption on molecular circadian rhythms within hippocampus has not been examined Time-restricted eating (TRE) in humans and time-restricted feeding (TRF) in animals, during which time of access to food is restricted to hours of the active phase, without calorie restriction, has emerged as an alternative strategy to protect against obesity and dysmetabolism. Human studies of TRE showed weight and fat loss as well as improved metabolic parameters in overweight and obese adult patients [20,21]. In animal models, re-alignment of food intake onto circadian rhythms by TRF restored the oscillations of liver transcripts related to metabolism and improved metabolic health [22–24]. In models of Alzheimer disease, TRF was also beneficial restoring part of the hippocampus circadian transcriptome and improving memory [25]. A recent study performed in mice exposed to 18 weeks of HFS diet at adulthood found that 1-week TRF protocol rescued spatial working memory and hippocampal plasticity but molecular rhythms were not examined [26]. The beneficial effect of TRF on juvenile HFS diet-induced long-term memory and hippocampal circadian transcriptome remains to be evaluated.

To fill these gaps, we identified behavioral and hippocampal molecular circadian disruptions in our model of juvenile consumption of HFS diet. Circadian rhythms of food intake, respiratory exchange ratio and energy expenditure but not of locomotor activity were altered in mice consuming HFS diet since weaning. The alteration of circadian metabolic rhythms was prevented by a 4-week TRF protocol. Therefore, we evaluated the effects of TRF in the memory impairments previously observed in HFS mice. TRF prevented the alteration in several hippocampal-dependent memory deficits observed in mice with unlimited access to HFS. TRF also normalized part of the hippocampal translatome altered by HFS diet and restored the circadian expression of genes that displayed blunted expression under *ad libitum* HFS diet. In particular, *ad libitum* HFS diet led to a reduced response of different genes of the thyroid hormone signaling pathway during memory formation. Remarkably, such alteration was fully rescued by TRF. Moreover, TRF restored the diurnal expression variation of a number of thyroid regulated genes involved in astrocyte maturation and glutamate reuptake. Finally, the intra-hippocampal administration of T3, the active form of thyroid hormone, rescued long-term object recognition memory deficits. Altogether, our results show that TRF prevents hippocampal-related memory deficits and establish an important role of thyroid hormones in HFS diet-induced memory impairments.

## Results

### Juvenile ad lib HFS diet induces, and TRF prevents, alteration of several circadian metabolic parameters

We have compared four groups of male C57Bl/6J mice that were either under a normal chow (NC) or HFS diet (45 kcal fat, 17% sucrose) for 12 weeks since weaning, provided *ad libitum* (ad lib) for the entire 12 weeks (NC ad lib, HFS ad lib) or ad lib for the first 8 weeks followed by time-restricted feeding (TRF) for the last four weeks (NC TRF, HFS TRF). First we showed that TRF significantly reduced the weight gain of the mice under HFS diet at week 11 (p=0.0133) and week 12 (p=0.0003) but did not have an impact on the weight of NC-fed mice (Fig. 1A, 3-way ANOVA, interaction Diet x TRF x Time: F_(12,3349)_=10.78, p<0.0001). Accordingly, in HFS mice, TRF was found to significantly reduce fat mass gain over 4 weeks (week 12-week 8), 7.82% vs. 12.04% respectively, for HFS TRF and HFS ad lib, p=0.0001 (Fig.1B, 2-way ANOVA, interaction Diet x TRF F_(1,50)_=4.577, p=0.0373). On the other hand, the fat mass gain did not significantly differ between NC TRF and NC ad lib (Fig. 1B).

**Fig. 1:**
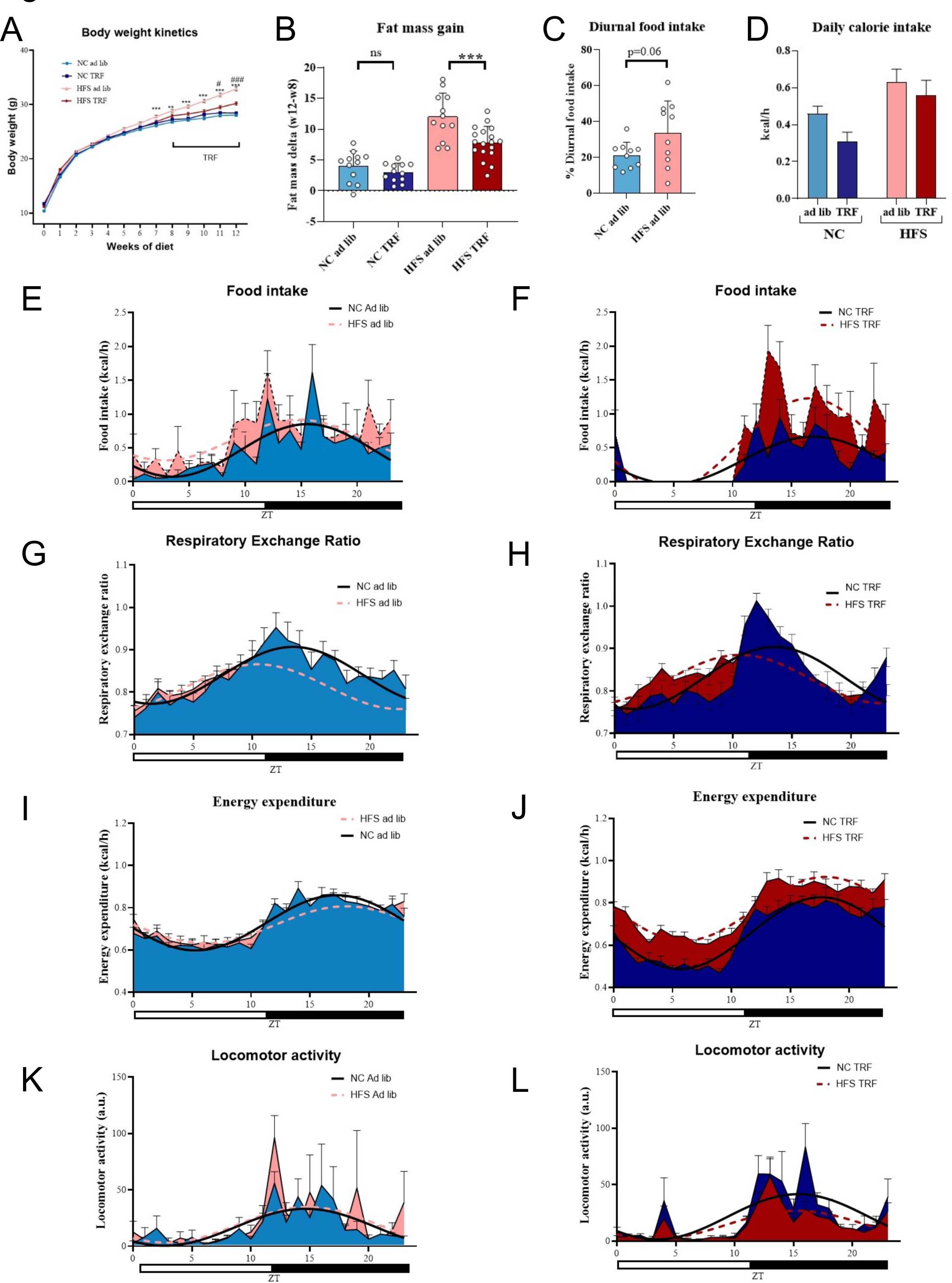
Juvenile ad lib HFS diet induces, and TRF prevents, alteration of circadian rhythms of food intake, respiratory exchange ratio and energy expenditure but not of locomotor activity. **A**: Kinetics of body weight across the 12 weeks of exposure; data from all the mice used in the study were compiled, n= 44-92 per group (some cohorts not weighted every week). Data were analyzed by 3-way ANOVA (Diet, TRF and Time (weeks of diet) factors, repeated measure for Time). **B**: Fat mass gain between week 8 and week 12 (measured by EchoMRI), unpaired t-test; n=12 per group except HFS TRF n=18. **E-M**: Metabolic rhythms measured by indirect calorimetry, on the left under ad lib and on the right under TRF regimen; n=10 per group. Mean (mesor), amplitude and acrophase values were calculated by Cosinor regression for each group. Data were compared by 2-way ANOVA analyses (Diet and TRF factors).

Using indirect calorimetry, we examined food intake, respiratory exchange ratio (RER), energy expenditure (EE), and locomotor activity rhythmicity during 24 hours in light-dark conditions. Cosinor regressions were used to calculate mesor (hourly mean levels), amplitude (difference between highest and lowest values), and acrophase (peak value) for each metabolic parameter. We found higher mean calorie intake in HFS than NC groups as expected (main effect of diet on food intake mesor F_(1, 36)_ = 12.30; p=0.0012), higher food intake amplitude in HFS TRF compared to both HFS ad lib and NC TRF (diet x TRF interaction on food intake amplitude F_(1, 36)_ = 6.171, p=0.0178) and an earlier food intake acrophase in ad lib than TRF animals (main effect of TRF on food intake acrophase, F_(1, 36)_ = 4.390; p=0.0433)(Fig. 1E-F). From the food intake measurements, we were able to calculate the diurnal percentage of food intake which was 33 ± 5% in HFS ad lib mice versus 21 ± 2% in NC *ad lib* mice (p=0.06, unpaired t test) (Fig.1C). Total 24h calorie intake was significantly impacted by HFS but not by TRF, (2-way ANOVA, main diet effect F_(1, 36)_ =11.45; p=0.0017) (Fig. 1D).

For RER, whose decreased and increased values reflect, respectively, higher utilization of lipids and carbohydrates, hourly mean RER was significantly lower in HFS ad lib than both NC ad lib (p<0.001) and HFS TRF (p=0.042) and there was no differences between HFS TRF and NC TRF (p=0.71) (diet x TRF interaction on mesor, F_(1,36)_= 5.905, p = 0.0202). Then, the acrophase was shown to be earlier in HFS than NC, and the amplitude was lower in HFS than NC groups but no effect of TRF was detected (main effects of diet on RER acrophase (F_(1, 36)_ = 54.55; p<0.0001 and RER amplitude, F_(1, 36)_ = 4.680; p=0.0372) (Fig.1G-H).

As for EE, we found higher levels of mean EE in HFS TRF mice than both HFS ad lib mice (p<0.0001) and NC TRF mice (p<0.0001) (diet x TRF interaction mesor, F_(1, 36)_= 78.14, p<0.0001). When looking at the EE amplitude, NC mice showed a higher amplitude than their HFS counterparts, and TRF animals had a higher EE amplitude than ad lib mice (main effects of diet, F_(1, 36)_= 9.723, P=0.0036 and TRF, F_(1, 36)_= 26.49, P<0.0001). There was no significant interaction nor main effects of diet and TRF on EE acrophase (Fig.1I-J).

Finally, there was no significant interaction nor main effect of diet and TRF on locomotor activity mesor, amplitude or acrophase (Fig.1K-L).

In sum, TRF on HFS diet, by resynchronizing food intake to day/night cycles, restored food intake amplitude, mean RER, mean EE but had no effect on locomotor activity.

### Juvenile ad lib HFS diet induces memory deficits that are prevented by TRF

Since TRF re-aligned metabolic rhythms onto circadian rhythms in mice under HFS diet, we examined whether the hippocampal-dependent memory impairments previously observed in HFS ad lib mice [10,13] were prevented by TRF. Indeed, TRF prevented the impairments on short-term object location memory (OLM) test (one-sample t test against 50%: p<0.001 for all groups except HFS ad lib; 2-way ANOVA interaction diet x TRF p<0.0001, HFS ad lib p<0.0001 compared to all other groups, Fig.2A). TRF prevented alterations on long-term object recognition memory (ORM) in HFS mice, tested in the morning, i.e. light phase (one-sample t test against 50%: p<0.05 for all groups except HFS ad lib and NC TRF; 2-way ANOVA: interaction p=0.0019, HFS ad lib p<0.004 compared to all other groups except NC TRF (Fig.2B). TRF also prevented impairments of ORM in HFS mice when tested in the evening (dark phase) (one-sample t test against 50%: p<0.05 for all groups except HFS ad lib, 2-way ANOVA, main diet effect p=0.033, HFS p<0.03 compared to all other group except p=0.06 compare to HFS TRF), (Fig. 2C). Regarding contextual fear conditioning, another hippocampal-dependent memory test, the impairments observed in HFS ad lib mice were only partially rescued by TRF since HFS TRF mice were not significantly different from neither NC ad lib nor HFS ad lib mice (2-way ANOVA, main diet effect, p=0.0011; NC ad lib or NC TRF vs. HFS ad lib, p<0.003 but vs HFS TRF p=ns; HFS ad lib vs HFS TRF p=ns) (Fig.2D). Notably, anxiety-related behaviors were unchanged in the four experimental groups whether tested in the elevated plus maze or in the light/dark box (2-way ANOVA, p=ns) (Fig. 2E &2F).

**Fig. 2:**
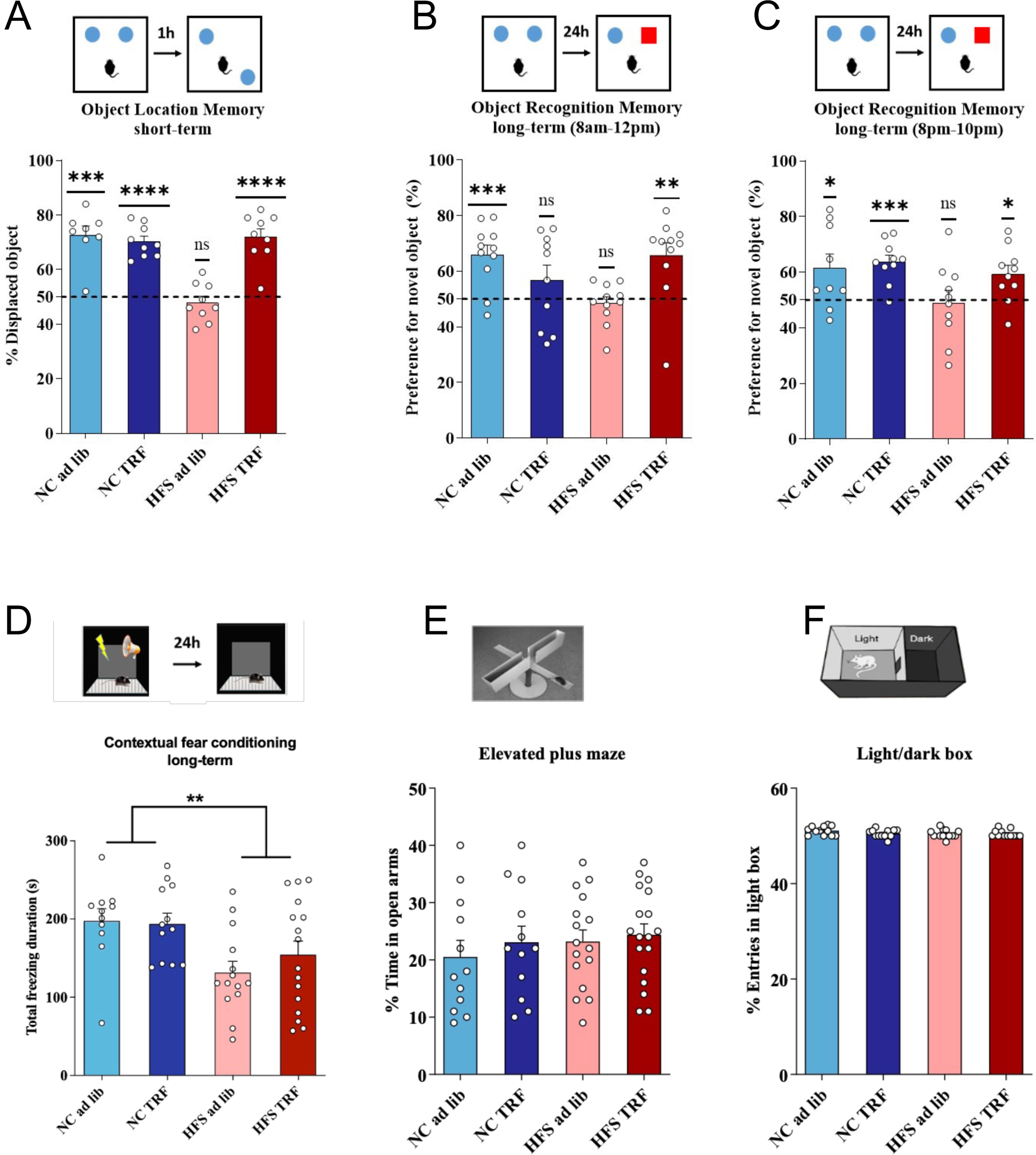
Juvenile ad lib HFS diet induces memory deficits that are prevented by TRF. **A**: Object Location Memory test, n=8-9 /per group; **B-C**: Object Recognition Memory test in the morning (n=10-11 per group) and the evening, n=9-10 per group. **D**: Contextual Fear Conditioning, n=12-16 per group. **E**: Elevated Plus Maze n=12-18 per group. **F**: Light/Dark box n=12per group. For OLM and ORM, each experimental group was assessed for a value of preference, analyzed by a one-sample t test between the group mean and 50%, that is, chance level as well as by 2-way ANOVA (Diet and TRF factors). CFC, EPM and LD data were analyzed by 2-way ANOVA (Diet and TRF factors). ns: p>0.05; *: p<0.05; **: p<0.01; ***: p<0.001; ****: p<0.0001)

### HFS ad lib diet impacts memory-induced hippocampal translatome that is partially rescued by TRF

In order to get insights into the mechanisms involved in HFS-induced memory impairments and their prevention by TRF, we examined the hippocampal molecular changes that accompanied the behavioral effects using a translatomic approach. Given that only a fraction of cells is activated in the hippocampus during memory training and consolidation, we employed a Translating Ribosome Affinity Purification technique to immunoprecipitate mRNAs currently being transcribed, using an antibody targeting the terminal phosphorylation sites (Ser244/247) of ribosomal protein S6 (pS6-TRAP) as in [27,28] followed by RNA sequencing to quantify mRNAs. The design of the pS6 TRAP-RNAseq experiments is depicted in Fig.3A. In a first experiment, mice from NC ad lib, HFS ad lib, and HFS TRF groups were each split into two-subgroup, one being euthanized under home cage conditions at zeitgeiber time (zt) 3 (HC zt3) and the other half culled 1 hour after a 10-min training of the ORM task that occurred between zt2 and zt4 (ORM+1h) to study the early phase of memory-induced genes. In a second experiment, mice from NC ad lib, HFS ad lib, and HFS TRF groups, were euthanized either under home cage conditions at zt15 (HC zt15) for half of each group or 12 hours (ORM+12h, around zt14-zt16) after a 10-min training of the ORM task that occurred between zt2 and zt4 for the other half of each group, to test the late phase of memory-induced genes as well as the diurnal expression variation by comparing HC zt15 vs. HC zt3 data and ORM+12h vs. ORM+1h. The group NC TRF was not analyzed because both metabolic rhythms and behaviors were similar to NC ad lib group. Tables of each differential analysis for each diet, as well as Enrichment analyses tables are available in supplementary data.

**Fig. 3:**
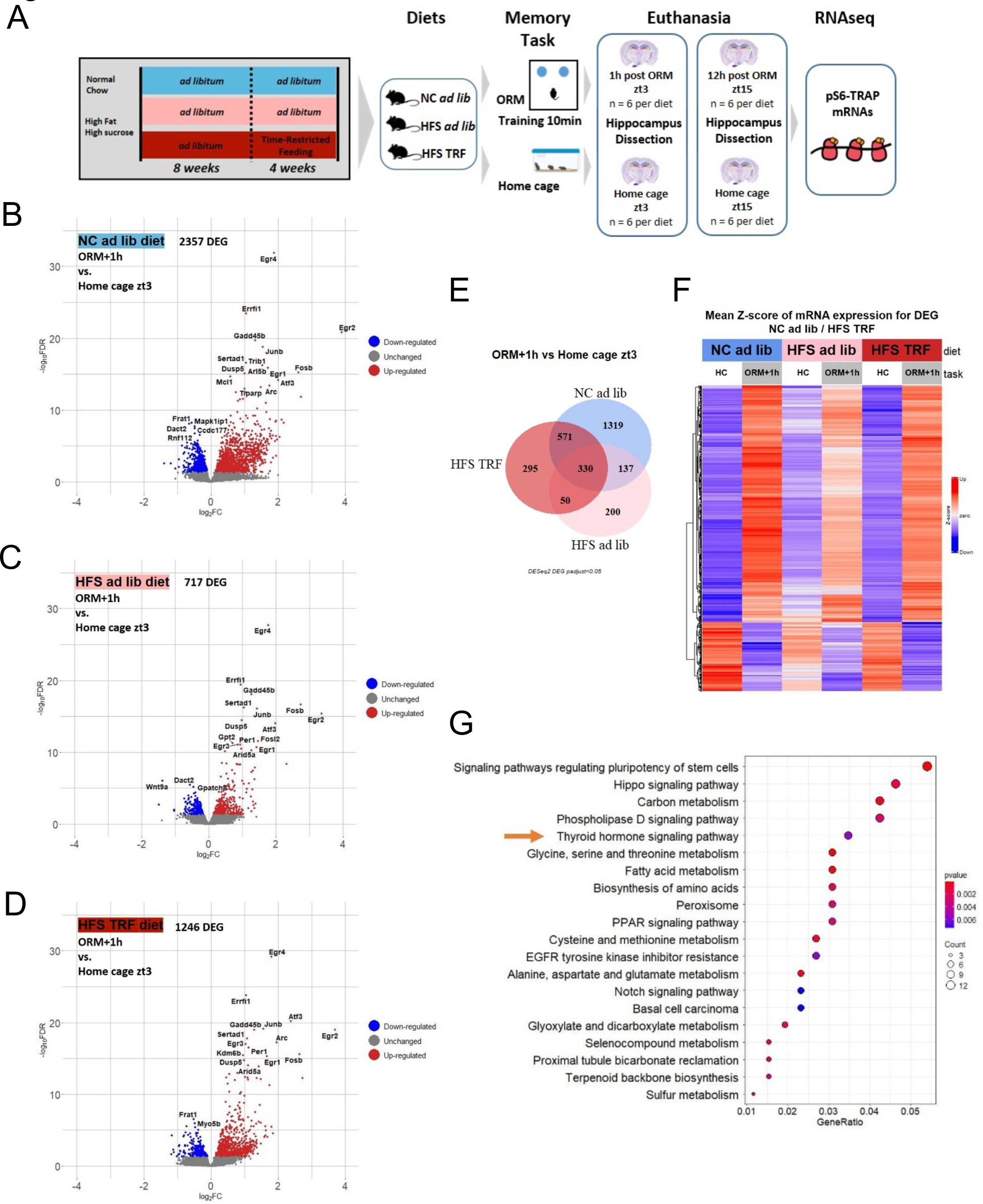
Juvenile ad lib HFS diet modulates circadian hippocampal translatome that is partially prevented by TRF. **A**: Experimental design. **B-D**: Volcano plots of the differential gene expression analyses in each diet comparing the early transcriptional response to memory (ORM+1h vs. HC zt3). **E**: Venn diagram of the differential gene expressed in each diet (ORM+1h vs. HC zt3). **F**: heatmap of mean z scores of the DEG that are shared between NC ad lib and HFS TRF groups. **G**: Enrichment analysis of the DEG that are shared between NC ad lib and HFS TRF groups. Differential gene expression analyses were done with DEseq2 package using a FDR p<0.05.

The data of all the RNAseq experiments were analyzed in a single statistical model using a false discovery rate (FDR) of p<0.05. As illustrated in the Volcano plots of the differential gene expression analyses comparing the conditions ORM+1h and HC zt3 of each diet (Fig. 3B-D), there were more genes up-regulated than down-regulated in each diet and the highest number of differentially expressed genes (DEG) was found in the NC ad lib group (2357), the lowest in HFS ad lib group (717) with HFS TRF mice displaying an intermediary number of DEG (1246). Interestingly, the analysis revealed a total of 571 DEG being commonly expressed in both NC ad lib and HFS TRF mice, while these genes were unchanged in HFS ad lib after ORM training (Fig. 3E). This subset of genes is of particular interest since it may be associated with the biological functions that are rescued by TRF to ameliorate HFS-induced memory deficits. Their mRNA levels, expressed as mean z scores, are illustrated in a heatmap (Fig. 3F). Enrichment analysis of these 571 DEG (Fig. 3G) revealed the involvement of these DEG in cell proliferation (pathways regulating pluripotency of stem cells, Hippo and Notch signaling), fatty acid (Fatty acid metabolism, Peroxysome, PPAR signaling) and amino acid metabolism (Biosynthesis of amino acid; Glycine, serine and threonine metabolism; Cysteine and methionine metabolism; Alanine, aspartate and glutamate metabolism). Among the top-enriched pathways, thyroid hormone signaling caught our attention due to their known involvement in memory processes [29]. Additionally, given that mean energy expenditure was lower in HFS ad lib compared to NC ad lib mice and enhanced in HFS TRF mice (Fig.1H), we hypothesized that circulating thyroid hormones may have been dysregulated, impacting both energy expenditure and memory through respectively their action on adipose tissue/ hypothalamus ([30]) and hippocampus ([29]).

When we examined the late-phase translational response of gene expression by comparing the condition 12 hours after the ORM task (ORM+12) to the home cage control at a similar time point (HC zt15), only 18 DEG were found rescued by TRF (Fig. S1), suggesting that the reprogramming of hippocampal translatome by the HFS diet takes place mainly in the early phase of memory consolidation. Most of these 18 genes are reported to be involved in neuronal plasticity, through neurogenesis or neurite outgrowth. None of these 18 genes were differentially expressed in NC ad lib and HF TRF, one hour after ORM.

Having identified the broad pattern of translatome-wide changes across diet exposure and memory changes, we sought to resolve gene co-expression networks to identify translatome changes that could be critical in determining the effect of diet and TRF on memory consolidation in its early stages, namely 1 h after ORM. We therefore used a weighted gene co-expression network analysis (WGCNA) [31] as a systems biology approach to identify modules (clusters) of highly co-expressed genes from the RNAseq data (ORM+1h vs HC zt3), in each diet. All tables related to this analysis are available in supplementary data. WGCNA yielded 6 clusters (modules) of highly correlated transcripts from NC ad lib, 10 clusters from HFS ad lib and 6 clusters from HFS TRF RNAseq data, the “grey” module representing unassigned transcripts (Fig. 3H-J). We then correlated the level of expression of each module eigengene (summarizing expression of the genes included in each module) with ORM versus HC conditions. NC ad lib displayed one module (brown) and HFS ad lib presented 2 modules (green and pink) highly correlated with ORM (Fig. 4A-B). Strikingly, HFS TRF mice displayed 2 modules correlated with ORM, one (blue module) highly enriched in genes expressed in NC ad lib mice brown cluster (245/298, 82%) and the other one (green) enriched in genes expressed in the HFS ad lib most significant module (62/121, 51%; Fig.S2 for details). Enrichment analyses of the 245 genes overlapping NC and HFS TRF (Brown-Blue) modules is presented in Fig.4D. In addition to the pathways displayed in the enrichment of Fig. 3G, we observed an enrichment in nervous system development, glial cells differentiation and regulation of endothelial cell migration as well as positive regulation of intracellular signal transduction, L-glutamate transmembrane transport and the glutamatergic synapse.

**Fig. 4:**
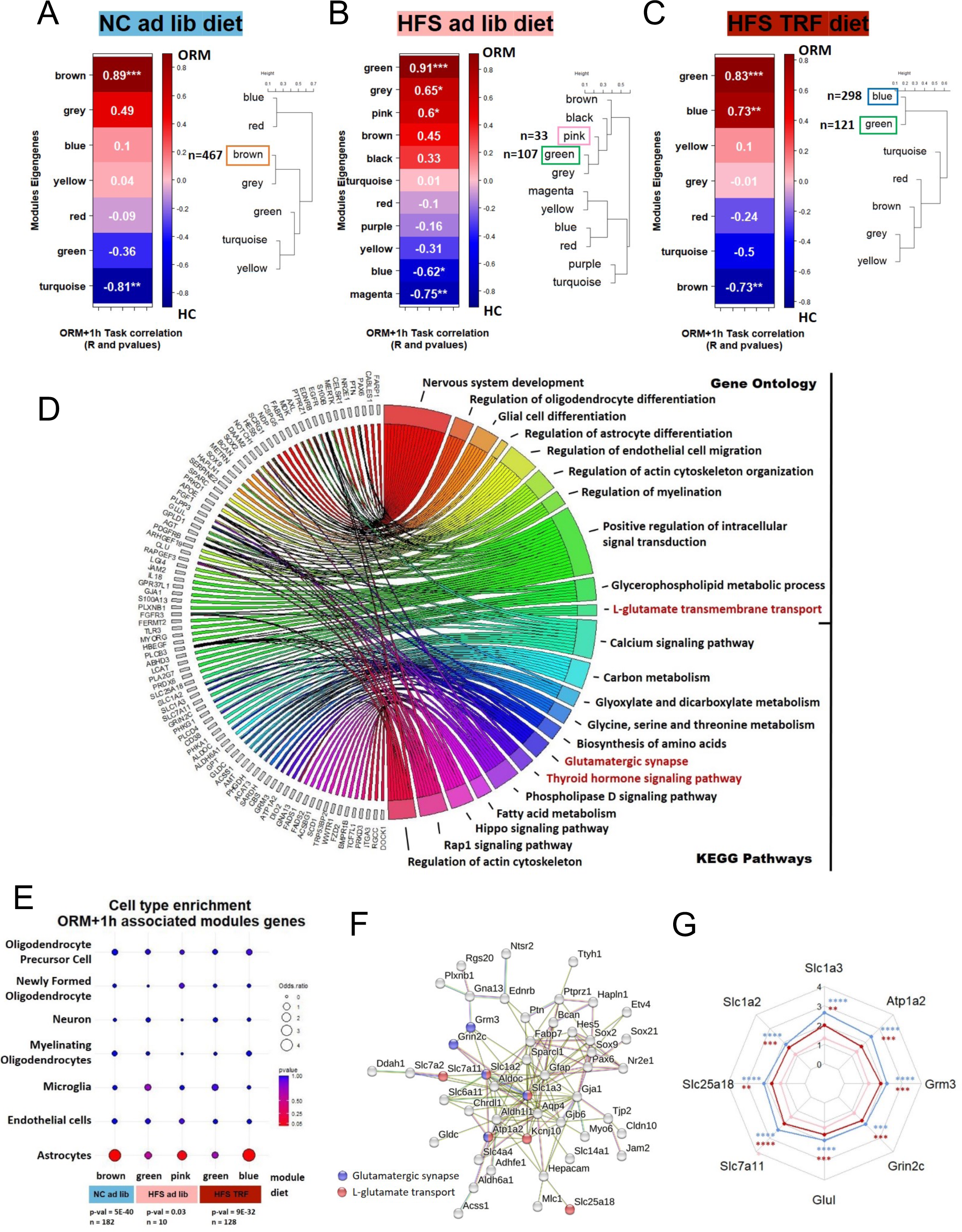
Co-regulated network analysis highlights astrocytic genes associated with memory. **A-C**: Modules of co-regulated genes in each diet as defined by WGCNA analyses. Each module is identified with a color name. The color scale on the right indicates the R values for the Pearson correlations (Module-trait correlation between eigengene expression in each module and ORM vs. home cage (HC). (P values for Pearson correlation are noted within each module). **D:** Enrichment analysis (Circos plot) of the genes that are correlated with ORM and in common between NC ad lib and HFS TRF modules. **E**: Cell-type analysis of the ORM-correlated genes for each module of each diet. **F**: Major network of the astrocytic genes in common between NC ad lib and HFS TRF modules (STRING software). **N**: Gene expression (RNAseq data) represented in a radar plot for genes involved in the regulation of glutamate neurotransmission. The stars shown on the top of each gene represent the p adjusted values from the DEseq2 global analysis.

Next, a cell type enrichment analysis [32,33] was performed on genes from each ORM-correlated modules in each diet. This analysis revealed that for mice under NC ad lib and HFS TRF, genes correlating with ORM where mostly expressed in astrocytes, which was not the case for mice under HFS ad lib as only 10 genes from the pink modules were expressed in astrocytes (Fig. 4E). We identified 123 astrocyte-expressed genes in common between NC ad lib and HFS TRF mice (Fig.S2) which were then processed in a gene network analysis. This analysis provided 5 clusters, of which the major one included 49 genes with several of them involved in the regulation of glutamate uptake and transmission (Fig.4F). These genes, namely GLAST (Slc1a3), GLT1 (Glutamate transporter 1, Slc1a2), GC2 (Glutamate/H(+) Symporter 2, Slc25a18), XCT (Cystine/Glutamate Transporter, Slc7a11), Na+/K+ ATPase pump (Atp1a2), glutamate metabolizing enzyme GS (Glutamine Synthase, Glul), NR2C (Glutamate Ionotropic Receptor NMDA Type Subunit 2C, Grin2c) and MGLUR3 (Glutamate Metabotropic Receptor 3, Grm3), contrary to NC ad lib and HFS TRF groups, are not significantly upregulated in mice under HFS ad lib diet (or less upregulated for XCT)), suggesting a deficiency in glutamate reuptake and recycling for mice under this diet (Fig.4G).

Collectively, the RNAseq analysis on memory-induced genes indicated that the hippocampal translatome is highly modified under HFS ad lib diet at the early phase of gene transcription. Part of the hippocampal translatome is rescued by TRF and includes thyroid hormone signaling as well as astrocytic genes involved in the regulation of glutamate neurotransmission.

### HFS ad lib diet impacts day-night variations of hippocampal translatome that is partially rescued by TRF

Since HFS ad lib diet is known to disturb molecular rhythms and TRF to reinstate them, we evaluated day-night gene expression by the comparison of DEG between first HC zt3 and HC zt15 (Fig. 5A-F) but also between ORM+12h (zt15) vs. ORM+1h (zt3) (Fig. 5G-L) in each diet. For the first comparison in home cage condition, a higher number of DEG was found in HFS TRF mice (2470) followed by NC ad lib (1735) and HFS ad lib (1391) as shown on the volcano plots and the Venn diagram (Fig. 5A-D). Importantly, a high number of genes (672) lost their day-night oscillation in HFS ad lib mice while this diurnal variation was rescued by TRF (Fig. 5D), as illustrated on the heatmap of mean z score mRNA expression (Fig. 5E). Interestingly, 19% (130/672) of these genes were part of the brown-blue gene modules correlated with ORM. Remarkably, enrichment analysis revealed that this subset of genes was enriched in thyroid hormone signaling and glutamergic synapse (Fig. 5F) pathways already identified in the ORM+1h vs HC zt3 analysis, hence indicating that rhythmicity of such pathways are of particular relevance for the regulation of memory processing. Lastly, the core clock genes Per3, Npas2 and Nr1d2 (REVERβ) showed a loss of day-night expression variation in HFS ad lib but not in the two other groups, further indicating the perturbation of hippocampal circadian rhythms in mice fed HFS ad lib (Fig. S3).

**Fig. 5:**
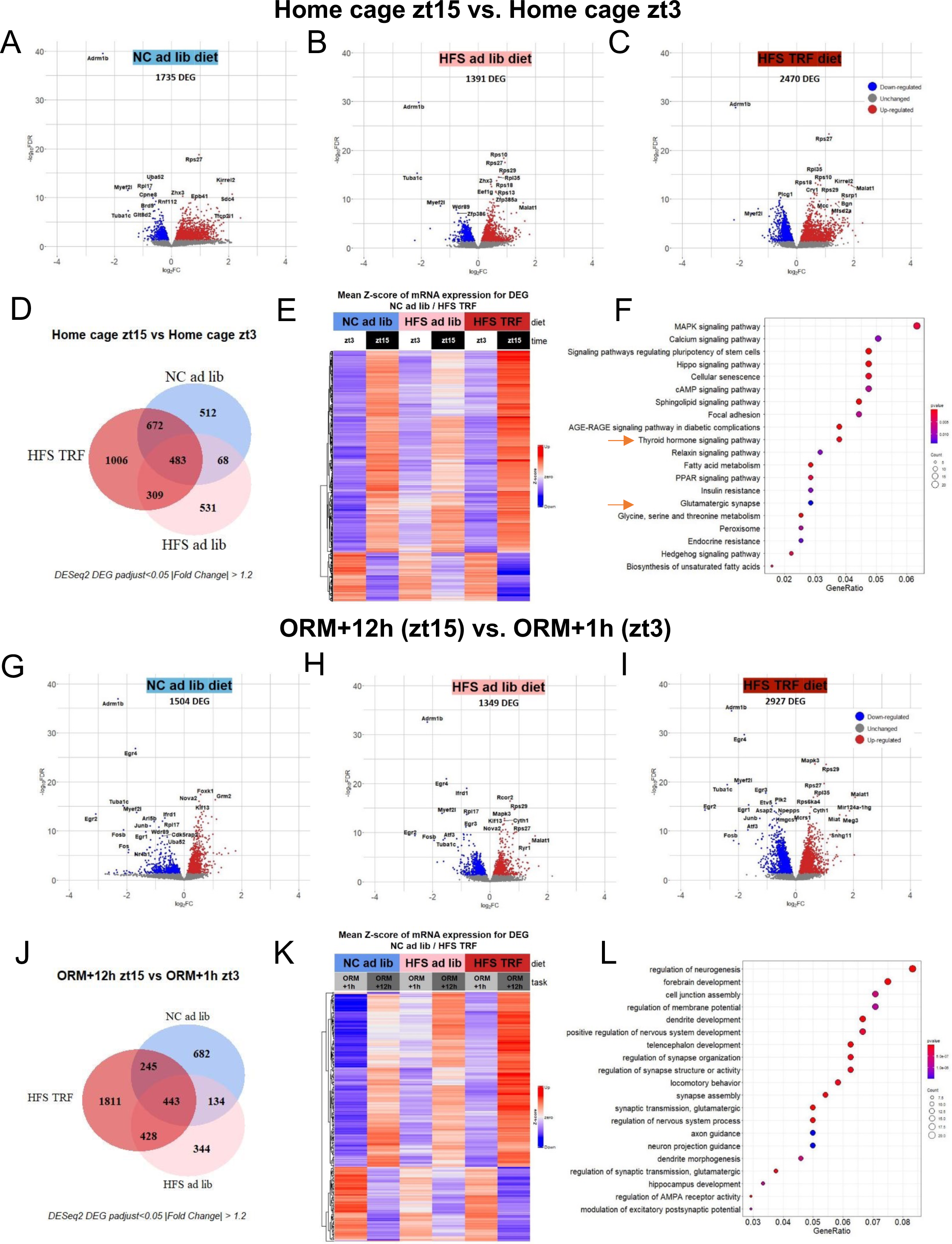
Juvenile HFS ad lib diet impacts day-night variations of hippocampal translatome that is partially rescued by TRF. **A-C**: Volcano plots of the differential gene expression analyses in each diet comparing diurnal variation of expression in home cage condition (HC zt15 vs. HC zt3). **D**: Venn diagram of the differential gene expressed in each diet (HC zt15 vs. HC zt3). **E**: heatmap of mean z scores of the DEG that are shared between NC ad lib and HFS TRF groups. Differential gene expression analyses were done with DEseq2 package using a FDR p<0.05 **F**: Enrichment analysis of the DEG that are shared between NC ad lib and HFS TRF groups. **G-I**: Volcano plots of the differential gene expression analyses in each diet comparing diurnal variation of expression after ORM (ORM+12h_ zt15 vs. ORM+1h_ zt3). **J**: Venn diagram of the differential gene expressed in each diet. **K**: heatmap of mean z scores of the DEG that are shared between NC ad lib and HFS TRF groups. Differential gene expression analyses were done with DEseq2 package using a FDR p<0.05 **L**: Enrichment analysis of the DEG that are shared between NC ad lib and HFS TRF groups.

In the day-night comparison ORM+12h vs. ORM+1h, the immediate early genes, markers of cell activity such as Fos, FosB, Egr1, Egr2, Egr4, were found down-regulated in each diet 12 hours after ORM compared to 1 hour (Fig.5G-I). As for home cage conditions, a higher number of DEG was found in HFS TRF (2927), followed by NC ad lib (1504) and HFS ad lib (1349) but only 245 genes were shared by HFS TRF and NC ad lib (Fig.5J), illustrated on the heatmap of mean z score mRNA expression (Fig. 5K). Moreover, the set of oscillating genes after ORM was different from home cage ones as only 15/245 (6%) genes were found in common for the NC ad lib/HFS TRF overlap. This gene subset was unrelated to ORM-correlated genes and to the genes induced 12 hours after ORM (ORM+12 vs. HC ZT15 DEG). Enrichment analysis for these post-ORM oscillating genes highlighted the implication of forebrain development, neurogenesis and dendrite development as well as synapse organization, structure, assembly and activity, and the glutamatergic synapse with the regulation of AMPA receptor activity (Fig.5L).

Overall, these results demonstrate that a large fraction of genes associated with memory function, and perturbed in HFS ad lib mice, exhibit baseline time-of day-specific expression, and indicate a central role of thyroid hormones signaling and glutamatergic synapse. In addition, genes that have lost diurnal variation of expression after ORM are also genes with important functions for memory formation. These data suggest that circadian dysfunction may impact the hippocampal translatome of HFS ad lib mice that is reinstated by TRF.

### Thyroid hormone signaling pathway is altered by HFS and partially rescued by TRF

Based on the aforementioned results, we decided to focus on thyroid hormone signaling pathway and to validate our translatomic analysis. First, we identified that Dio3, encoding iodothyronine deiodinase 3 enzyme, which metabolizes the active thyroid hormone T3 in inactive reverse T3 (rT3), was the most down-regulated gene, 1 hour after ORM, in NC ad lib and HFS TRF mice while its expression was unchanged in HFS ad lib mice. Dio2, encoding for iodothyronine deiodinase 2 enzyme, which converts T4 to active T3, is highly induced by the memory task in all groups, although with a lower fold change in HFS ad lib (Fig.S4). Consequently, the ratio of Dio2/Dio3 hippocampal expression revealed a decreased ratio in HFS ad lib mice, suggesting a reduced T3 availability within the hippocampus of these mice that is rescued by TRF (Fig.6A). The thyroid transporters MCT8 (Slc16a2) (Fig.6B) and OATP1 (Slco1c1) gene expression (Fig.6C) was found unchanged in all diet. Thyroid hormones have two receptors THRα and THRβ, each of them displaying two isoforms. Thrb2 but not Thrb1 isoform expression was detected in hippocampus by the RNAseq analysis, and was found unchanged after ORM in all diet (Fig.6D). As for THRα, both Thra1 and Thra2 were highly expressed in the hippocampus. Thra1 expression was not modified by ORM nor diet while the expression of Thra2 was significantly changed by ORM in NC ad lib and HFS TRF mice (Fig.S3). The ratio of the two isoforms increased after ORM in NC ad lib and HFS TRF but not in HFS ad lib (Fig. 6E). Since Thra2 is a dominant-negative isoform that inhibits the transcriptional activity of Thra1, the reduced ratio of Thra1/Thra2 in mice under HFS ad lib diet indicates that 1 hour after ORM, THRα1 signaling may be less efficient in these mice due to the increased inhibitory action of THRα2. Because THRα1 and THRα2 isoforms are generated by the differential splicing of the Thra gene, we examined differential exons usage on the RNAseq data of each diet. The DEXseq analysis revealed that a THRα1-specific exon is differentially used in mice under NC ad lib (p=0.037) and HFS TRF (p=0.030) after ORM but not in mice under HFS ad lib diet (Fig.6F). This result is in agreement with the reduced ratio of Thra1/Thra2 expression and a lower THRα1 signaling in HFS ad lib mice since 1h after ORM the Thra1 isoform is favored in NC ad lib and HFS TRF but not in HFS ad lib mice. Such locally fine-tuned mechanism of THRα regulation has never been described before to our knowledge.

**Fig. 6:**
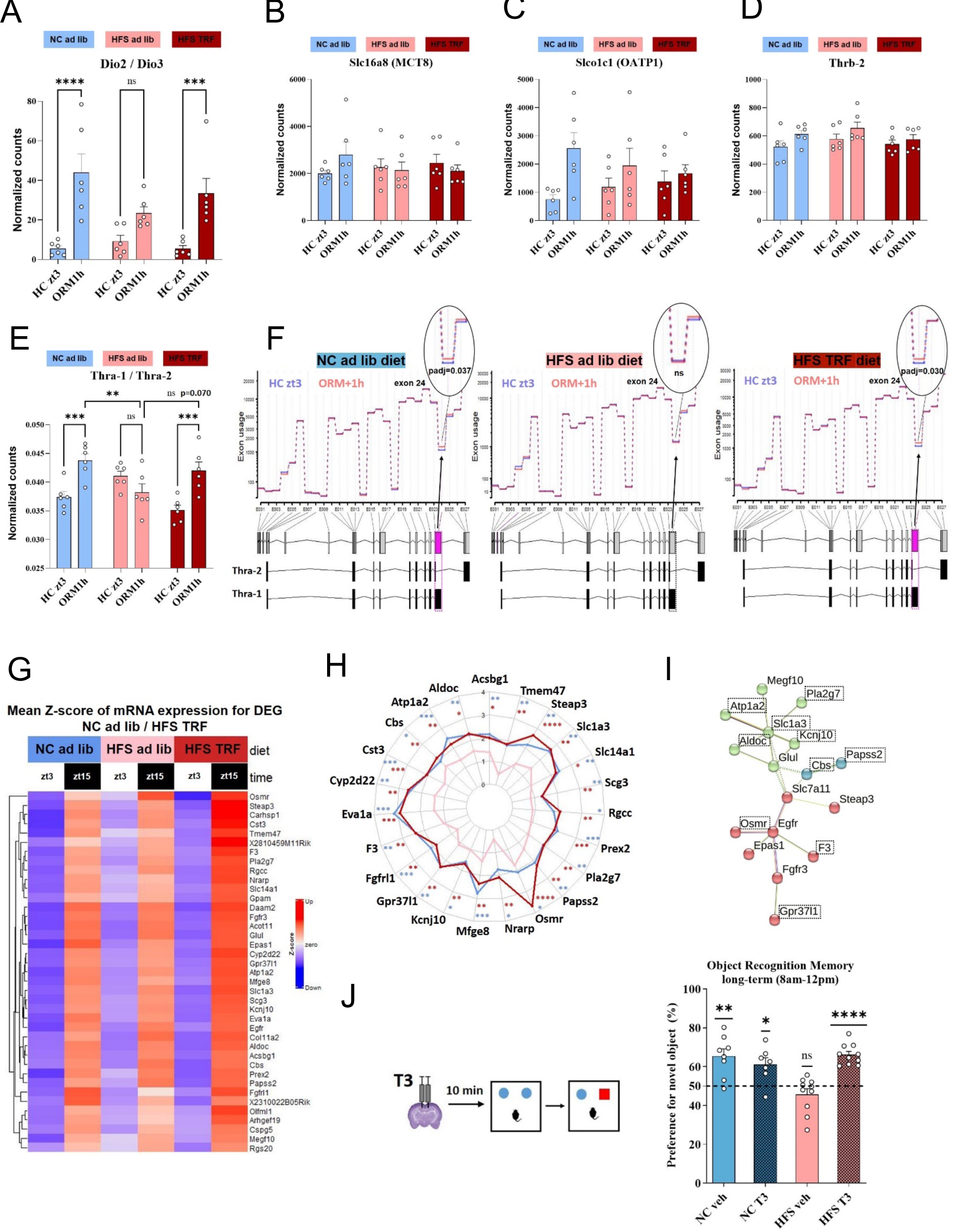
Thyroid hormone signaling pathway contributes to HFS diet-induced memory impairments and TRF prevention. **A**: Gene expression ratio of Dio/Dio3 (normalized counts from DEseq2 analysis). **B**-**-D**: Slc16a8, Slco1a1 and Thrb2 gene expression from the DEseq2 analysis **E**: Ratio of Thra1/Thra2 from the Deseq2 analysis. **F**: DEXseq analysis for THra gene in each diet. The differentially used exon is highlighted in pink and the p adjusted value from the DEXseq analysis indicated for each diet. **G**: Heatmap of mean z scores of DEG from genes that belong to the ORM-correlated gene network and target genes of thyroid hormones, in common between NC ad lib and HFS TRF. **H**: Radar plot of genes from **G** that display day-night variation of expression. **I:** Network constructed from genes described in **G**. The genes framed with a dooted line have a diurnal variation of expression that is lost in HFS ad lib mice. **J:** Long-term Object Recognition Memory (ORM) results of NC ad lib and HFS ad lib mice infused with either vehicle or T3 within hippocampus, n=7-10 mice per group, each experimental group was assessed for a value of preference, analyzed by a one-sample t test between the group mean and 50%, that is, chance level as well as 2-way ANOVA (Diet x T3 treatment). ns: p>0.05; *: p<0.05; **: p<0.01; ***: p<0.001; ****: p<0.0001)

We further looked at the enrichment of the early phase-induced memory genes (DEG ORM+1h vs zt3), for thyroid hormone signaling in each diet by comparing the list of DEG with a mouse brain atlas of thyroid hormones’ target genes recently published [34]. We found that in NC ad lib mice, 277 of the DEG are thyroid hormone targets, 72 in HFS ad lib and 155 in HFS TRF. We then focused on the thyroid hormones target genes that intersect between NC ad lib and HFS TRF groups (85 genes). We found that 46% (39/85) of them belonged to the ORM-correlated genes (brown and blue modules overlap of Fig.4). The mean z scores of mRNA expression of these 39 genes is shown in a heatmap (Fig.6G). We then found that 24 out of the 39 genes lost their diurnal expression variation under HFS ad lib diet by looking at their expression at zt15 vs. zt3 in home cage condition (Fig.6H). Among the 39 genes, 17 were linked in a network that underscored their implication in the glutamate transport and metabolism and astrocyte differentiation (Fig.6I).

### Thyroid hormone infusion within the hippocampus rescues memory in HFS ad lib mice

Given the convergent lines of evidence pointing towards the involvement of thyroid hormones in the impaired memory of HFS ad lib mice, we went on by assessing the functional impact of thyroid hormone on hippocampal-dependent memory processes in NC ad lib and HFS ad lib groups. For this, T3 hormone was locally and bilaterally infused within the dorsal hippocampus (1.2ng/0.3μl/side) 10 min before ORM training. This hippocampal T3 infusion rescued long-term ORM deficit in HFS ad lib mice without affecting performance of NC ad lib mice ((one-sample t test against 50%: p<0.02 for all groups except HFS ad lib; 2-way ANOVA interaction diet x TRF p<0.001, HFS ad lib p<0.01 compared to all other groups, Fig. 6J) demonstrating that deficient thyroid hormones action in hippocampus plays an important role in the memory impairments of HFS ad lib mice.

## Discussion

TRF without calorie restriction is currently the favored type of intermittent fasting strategy to fight obesity and its comorbidities because of its simplicity of implementation in human settings and the valuable effects observed in both animal models and humans. Studies analyzing the improvement of brain health and disease through TRF are only emerging [25,26,35]. The present study contributes to this effort by showing that a 4-week TRF protocol is salutary for short and long-term memories that are altered in mice fed HFS diet *ad libitum* during peri-adolescence. We provide evidence that thyroid hormone signaling contributes to the memory impairments induced by the HFS diet and their rescue by TRF, through modulation of astrocytic genes involved in the regulation of glutamate transmission.

First, we observed that mice under 12 weeks of *ad libitum* HFS diet display a clear tendency to eat more during the inactive light phase (zt0-zt12) than do the mice under NC diet as found by others [14,15,36,37]. Under TRF the HFS fed mice still present a higher food intake than NC mice but importantly feeding is now aligned onto day/night cycles. Increased energy expenditure seems responsible for the reduced fat mass gain in HFS TRF mice compared to HFS ad lib mice since locomotor activity was found unchanged and total food intake was not reduced by TRF in HFS mice (Fig.1). These effects of TRF on metabolism, especially the critical role on energy expenditure, have been observed in other studies [23,36] but not always [26,38]. These discrepancies may be due to varying diet composition, duration and age of exposure onset to the HFS diet and also by different schedule and length of TRF across studies.

In addition to its beneficial actions on metabolism, we found that TRF prevented both short-term object location and long-term object recognition memory deficits in our model of juvenile consumption of HFS diet. In our hands, the alteration of contextual fear conditioning observed in mice under ad lib HFS was only partially rescued by TRF, likely due to the complexity of the memory task (Fig.2). In a model of 18 weeks of HFS diet exposure at adulthood, spatial working memory (spontaneous alteration in a T maze) was found altered and then rescued by a single week of TRF [26]. This spatial short-term memory test was not included in our experimental design but we also found that TRF can prevent short-term memory impairment (OLM data). TRF also rescues short and long-term memory in Alzheimer’s mouse models using both the ORM and the radial arm maze paradigms [25]. Thus, our data comfort the idea that TRF can have a positive action on short and long-term memory when altered in mice.

The formation of long-term memory is dependent on the activation of gene transcription and de novo protein synthesis [39]. Concerning the early phase of transcriptional response (ORM+1h vs. HC zt3), our RNAseq data confirm, in each diet, the up-regulation of activity dependent transcription factors such as Egr2, Fosb, Fos, Egr1, Arc, Nr4a1 and DNA methylation modifiers such as Gadd45b found to be important in neuronal activity during memory formation [40]. The gene Per1, involved in age-related cognitive decline was up-regulated in each diet [41]. Enrichment analyses of this set of rescued genes highlighted the implication of thyroid hormone signaling which caught our attention given that these hormones have a central role in hippocampus-dependent cognitive functions such as learning and memory [29,42,43]. Furthermore, these hormones are also critical for energy expenditure through thermogenesis by their action on adipocytes but also on hypothalamus [30]. Thus, a global dysregulation of thyroid hormones may explain both the lower energy expenditure and the impaired memory in HFS ad lib mice.

When we examined the hippocampal translatome 12 hours after the ORM, we detected only 18 genes that were rescued by TRF, demonstrating that TRF affects particularly the first transcriptional response of memory consolidation. Additional time points would be necessary to explore the dynamic changes in gene expression across time after object recognition learning.

We further delved into the RNAseq data of the early phase of transcription (ORM+1h vs. HC zt3) by investigating the presence of co-regulated gene networks, their correlation with memory formation and the cell types involved. These analyses indicated that co-regulated gene networks being associated with memory formation in NC ad lib and HFS TRF mice were expressed predominantly in astrocytes where they play a key role in regulating central nervous system development, synaptic plasticity and glutamate buffering. Remarkably, several astrocytic glutamate transporters, such as GLAST and GLT-1 had their hippocampal expression very significantly up-regulated in NC ad lib and HFS TRF but unchanged in HFS ad lib one hour after ORM. These transporters are crucial for glutamate clearance as they ensure the reuptake of glutamate from the synaptic cleft. Among the genes of the network were also Atp1a2 and ATp1b2 genes that are encoding subunits of Na^+^, K^+^-ATPase pumps, also present in astrocyte membrane, and necessary for the activity of the glutamate transporters [44]. The gene Glul was also present in the memory-associated network and encodes glutamine synthase (GS) which controlled the termination of glutamate transmission by metabolizing glutamate into glutamine within astrocytes. Furthermore, the metabotropic glutamate receptor MGLUR3 and the ionotropic glutamate receptor NR2C encoding respectively the genes Grm3 and Grin2c displayed the same pattern of mRNA expression than the glutamate transporters. The absence of up-regulation of all of these genes in HFS ad lib mice after ORM likely lead to alterations of the clearance of glutamate that could change tonic excitation and synaptic currents mediated by glutamate receptors [45]. Fascinatedly, a recent study [46] revealed that the glutamate transporters GLAST and GLT-1 were found down-regulated in the ventral hippocampus of mice fed with a high fat diet for 12 weeks and responsible for an increase in extracellular level of glutamate measured by real-time recording biosensors. Along the same lines, GLT-1 functioning was found impaired in the orbitofrontal cortex of rats fed a high-fat diet for 40 days, leading to an elevation of extra-synaptic glutamate [47]. Although not measured in this study, excessive glutamate levels are likely in our model. A previous study reported that GLAST and GLT-1 were up-regulated in mouse hippocampus (by 27% and 32% respectively) after one week of HFS diet together with increased glutamate uptake velocity but GS was down-regulated as well as NR2B glutamate receptor density [48]. Another study showed that MGLUR2/3 receptor blockade restored impaired LTP in rats fed HFS diet for 6 months [49]. These results look contradictory to ours but can be explained by the length of HFS exposure, very short-term in the Valladolid-Acebes’ report and very long-term in Karimi’s experiment. Indeed, when 2-3 months HFS diet duration was compared to 6-7 months in the same study, LTP was found increased and decreased respectively [12]. Also of great interest for our study, thyroid hormones were reported to regulate the pattern of astrocytes maturation in cerebellum through THRα1 [50] and to up-regulate GLAST and GLT-1 in order to protect astrocytes and neurons against glutamate toxicity [51].

Thanks to the RNASeq data obtained at two time points 12 hours apart (HC zt15 vs HC Zt3), we were able to identify the genes that loss day-night expression variation in HFS ad lib mice and that were restored by TRF. A large fraction of these genes belongs to the memory-associated network identified above, and are enriched for thyroid hormones signaling and glutamatergic synapse pathway. In addition, genes that have lost diurnal variation of expression after ORM (ORM+12h vs ORM+1h) are also genes with important functions for memory formation. Overall, these data suggest that circadian dysfunction may impact the hippocampal translatome of HFS ad lib mice that is reinstated by TRF.

Given the aforementioned data, we then decided to focus on thyroid hormone signaling and its potential role on the memory deficits observed in HFS ad lib mice and their restoration in HFS TRF mice. The differential gene expression analysis showed that Dio3 was the most down-regulated gene in NC ad lib and HFS TRF mice in the hippocampus, which led us to compare the Dio2/Dio3 mRNA ratio levels between diets, a measure considered as a good proxy to estimate local bioavailability of the active form of thyroid hormone, i.e. T3. The Dio2/Dio3 ratio increased considerably 1 hour after ORM in NC ad lib and TRF but not in HFS ad lib, suggesting a deficit of active T3 in the hippocampus of HFS ad lib mice.

As for THRα, the ratio of the two isoforms indicated that THRα1 signaling might be increased after ORM in both NC ad lib and HFS TRF, potentially due to the relief of THRα2 inhibition, which does not occur in HFS ad lib mice. This result was comforted by the finding of differential exon usage in favor to THRα1 after ORM in NC ad lib and HFS TRF groups but not in HFS ad lib mice. Whether the high levels in dominant-negative THRα2 isoform after ORM impinge THRα1 signaling during memory formation in hippocampus of HFS ad lib mice is thus possible and interesting but remains to be demonstrated. Interestingly, such fine-tuning of thyroid hormone local action has been described for a patient holding a specific genetic mutation in the THRα gene sequence, resulting in an increased THRα2 antagonism and leading to neuronal hypothyroidism and intellectual disability [52]. Additionally, THRα1 represents 70% of thyroid hormones receptor proteins in the brain and as mentioned above, it has a specific role in astrocyte maturation [50,53] and modulates GLAST and GLT-1 [51]. Furthermore, we were able to identify that 16% of the ORM-correlated genes in NC ad lib and HFS TRF mice are also direct target of thyroid hormones with most of them showing day-night variation. Thus, an alteration of THRα1 signaling in HFS ad lib mice is congruent with an impairment in the genes regulating astrocytes differentiation and glutamate reuptake.

Finally, we demonstrated the importance of deficient T3 action in the hippocampus of HFS ad lib mice, by showing that the infusion of T3 directly within hippocampus restored the long-term ORM of HFS ad lib mice.

In conclusion, this study demonstrates that male mice fed a HFS diet since weaning present deficiency in hippocampal T3 hormone availability and strongly suggest that THRα1 signaling is impaired, leading to long-term memory impairments, through dysregulation of genes implicated in astrocyte maturation and glutamate clearance. TRF restores a higher day-night expression variation of part of these dysregulated genes modulated by thyroid hormones, that may prevent the long-term HFS diet induced memory deficits. A limitation of this study is that females were not analyzed. Their metabolic response to HFS diet is different from males and they do not present impairment in contextual fear memory but do display alteration in long-term object recognition memory (N’Diyae et al, 2024, *under revision*). The beneficial effect of TRF and the importance of thyroid hormone in female mice fed HFS diet since weaning is ongoing. Time-restricted eating has proven to be efficient to ameliorate metabolic parameters in obese patients including adolescents [54,55]. Now, it will be interesting to measure the effects of time-restricting eating on memory performances in adolescent patients with obesity.

## METHODS

### Animals and Time-Restricted Feeding (TRF)

Male C57BL/6J mice aged 3 weeks (Janvier Labs, France) were randomly divided into groups of 6 per cage (45x 25x 20 cm, containing a cardboard house, nesting material and a small wooden stick) and had *ad libitum* access to a normal chow diet (NC; 2.9 kcal/g; 8% lipids, 19% proteins, 73% carbohydrates; A04, NEUTRAL) or a high-fat and sugar diet (HFS; 4.7 kcal/g; 45% lipids, 20% proteins, 35% carbohydrates of which 50% is sucrose; D12451, Research Diet). All animals were housed in a temperature-controlled room (22 ±1◦C) maintained under a 12h light/dark cycle (lights on at 7:30 am; Zeitgeber time (zt) 0) and had free access to food and water for 8 weeks. From week 8 to week 12, NC and HFS mice were divided in 4 groups (NC *ad lib* and HFS *ad lib* groups with unlimited food access; NC TRF and HFS TRF with time-restricted access to food from zt11 to zt1. Animal weighing was performed once per week for most cohorts, fat mass (in grams) was measured by nuclear magnetic resonance (NMR, minispec LF90 II, Bruker, Wissembourg, 67166) after 8 and 12 weeks of diet exposure in one cohort of mice. Behavioral procedures started after 12 weeks of diet exposure. All animal care and experimental procedures were in accordance with the INRAE Quality Reference System and French (Directive 87/148, Ministère de l’Agriculture et de la Pêche) and European legislations (Directive 86/609/EEC). They followed ethical protocols approved by the Region Aquitaine Veterinary Services (Direction Départementale de la Protection des Animaux, approval ID: B33-063-920) and by the local animal ethic committee of Bordeaux CEEA50 (APAFIS #22684 and #32843). Every effort was made to minimize pain and discomfort and reduce the number of animals used.

### Calorimetry and feeding patterns

Daily patterns of energy expenditure and respiratory exchange ratio were determined in individual metabolic cages using an open-circuit indirect calorimetry system (Addenfi, Les Cordeliers, Paris, France). Energy expenditure and respiratory exchange ratio, defined as the ratio of produced CO_2_ production and consumed O_2_, were calculated using AlabSuite (v1.55, Addenfi). This set-up also recorded feeding behavior using an automated weighing system of the feeder (detection threshold of 0.05 g), as well as total locomotor activity estimated by weighing sensors integrated into the bottom of the cage. Mice were acclimatized to the metabolic cages during 2 consecutive days. Measurements and analyses were performed during the third day.

### Behavioral procedures

Behavioral activity of each session was recorded through a camera connected to a monitor outside the experimental room allowing the experimenter to visualize the mice during the session and to evaluate and analyze the behaviors later on. Animals were handled every day (1-2 min) for 3 days before the beginning of each behavioral procedure.

### Object Recognition Memory (ORM)

This test was performed as described by [Leger 2013][56] in order to evaluate long-term novel object memory. For the training session, mice were placed into an open field arena (40cm x 40cm) containing two identical objects (cylinders) and were left to explore these called familiar objects until they reached a criterion of 20s of total exploration for both objects. Exploration time was registered when the snout of the mouse was directed towards the objects from a distance shorter than 2 cm, and climbing on the objects was not recorded as exploration. During the test session, performed either 24h (or 48h for T3 infusion experiment) after the training session, mice were placed in the same arena and exposed to two objects, one familiar (cylinder) and one novel object (lab glass bottle). The time exploring these objects was again quantified until a criterion of 20s of total exploration for both objects was reached. Thereafter, a discrimination index was calculated as the difference in time exploring the novel and familiar object, expressed as the ratio of total time spent exploring both objects. A maximum cutoff of 10 minutes was established for both sessions, and animals that explored the objects less than 12s were excluded from the analysis. Exploration time was manually quantified, by a trained experimenter outside the experimental room.

### Object Location Memory (OLM)

This test was performed in order to evaluate object location memory. For the training session, mice were placed into an open field arena (40cm x 40cm) containing two identical objects (cylinders) and were left to explore these called familiar objects until they reached a criterion of 20s of total exploration for both objects. Exploration time was registered when the snout of the mouse was directed towards the objects from a distance shorter than 2 cm, and climbing on the objects was not recorded as exploration. During the test session, performed 1h after the training session, mice were placed in the same arena and exposed to the same objects (cylinders) with one of them moved to the opposite angle of the arena. The time exploring these objects was again quantified until a criterion of 20s of total exploration for both objects was reached. Thereafter, a discrimination index was calculated as the difference in time exploring the moved object and the familiar object, expressed as the ratio of total time spent exploring both objects. A maximum cutoff of 10 minutes was established for both sessions, and animals that explored the objects less than 10s were excluded from the analysis. Exploration time was manually quantified, by a trained experimenter outside the experimental room.

### Contextual Fear Conditioning

An unpaired fear conditioning protocol was used as an alternate Pavlovian conditioning procedure capable of robustly producing contextual fear memory in mice. The day before fear conditioning, all mice were individually placed for 4 min into a white chamber (square, 20×20 cm) with an opaque PVC floor (neutral context). The box was cleaned with 1% acetic acid before each trial. The day after (Day 1), acquisition of fear conditioning was performed in a transparent enlightened conditioning chamber (square, 24×24 cm) (conditioning context). The box was cleaned with 70% ethanol before each trial. The floor of the chamber consisted of stainless-steel rods connected to a shock generator. During the unpaired conditioning, tones and foot-shocks were presented at random non-overlapping times over a total time of 232 seconds (s). More precisely, 100 s after being placed in the conditioning context, animals received a shock (0.3 mA, 1 s), then, after a 20 s interval, a tone (70 dB, 1000 Hz, 15 s) was delivered; finally, after a 30 s delay, the same tone then the same shock spaced by a 30 s interval were presented. After 20 s, animals were returned to their home cage. The next day, after a 24h delay, mice were submitted to two memory retention tests (Day 2). In the tone re-exposure test, mice were placed in the neutral familiar context during 6 minutes (min) divided in three successive sessions: one before (first 2 min), one during (next 2 min), and one after (2 last min) tone presentation. Two hours later, mice were submitted to the contextual re-exposure test: they were placed for 6 min in the conditioning context without any tone or foot-shock. Freezing behavior of animals, defined as a lack of all movement except for respiratory-related movements, was used as an index of conditioned fear response and continuously recorded for off-line s-by-s scoring of freezing by an observer blind to experimental groups. During training, freezing behavior was scored during the 20 s prior to the first foot-shock and the 20 s after each of the two foot-shocks to rule out any obvious signs of generalized fear, lethargy, or sickness that could confound exploratory behavior. During tests, to compare the freezing time in the neutral and conditioning contexts, data are presented as the percentage of time scored as freezing during the first 2 min in both contexts (i.e. before tone presentation in the neutral context).

### Light dark box test (LDT)

The LDT is a standardized test to assess anxiety in rodents. It is based on their innate aversion to brightly lit areas, and their spontaneous exploratory behavior when exposed to a novel environment. The light/dark box consisted in two compartments connected to each other by a small opening (width, 7 cm; height, 7 cm). The first compartment was made of white Perspex (length, width and height: 27 cm), illuminated by a white bulb delivering 300 lx. The second compartment, smaller than the first one (length: 18 cm; width and height: 27 cm), was made of black Perspex and illuminated by a red bulb delivering < 5 lx. Both white and red bulbs were located 37 cm above the apparatus floor. Each mouse was placed in the center of the white compartment facing the small opening. When the mouse entered into the black compartment, the number of transitions into the white compartment and the relative time spent therein were measured for 5 min.

### Elevated plus maze

The elevated plus-maze apparatus consisted of two opposing open arms (30 cm × 8 cm) and two opposing closed arms (30 cm × 8 cm × 15 cm) extending from a 8 cm × 8 cm central platform and elevated 1 m from the ground. Each mouse was first placed onto the central platform facing an open arm and then left to freely explore the apparatus for 5 min. The number of entries in the open or closed arms, as well as the time spent on the various sections of the maze, was recorded. The percentage of time spent in the open arms ((time in the open arms) / (time in the open + closed arms) × 100) was used as an index of anxiety, whereas total number of entries in the closed arms was used as an index of locomotor activity.

### T3 intra-hippocampal infusion

After 10 weeks under NC (n=15) or HFS (n=20), mice were anesthetized with air/isoflurane (4.5% induction; 1.5% maintenance) at 1L/min, injected with the analgesic buprenorphine (Buprecare, 0.1 mg/kg s.c. in 0.9% saline) and the non-steroidal anti-inflammatory drug carproxifen (Carprofene, 20mg/kg s.c. in 0.9% saline) and were placed into the stereotaxic apparatus (David Kopf Instruments). The scalp was shaved, disinfected with 10% povidone iodine and locally anaesthetized with a subcutaneous injection of lidocaine (Lurocaine 20mg/ml, 0.1ml non-diluted) and a midline incision made over the top of the skull. Bregma and lambda were located and marked to determine implant position. Bilateral implant (8 mm stainless steel guide cannula) coordinates were 2mm posterior posterior and 1mm ventral to Bregma and 1.3mm distal from the midline on both sides targeting dorsal CA1. Guide cannulae were secured in place with dental cement. Mice were kept on a heating pad until recovery. Mice were single housed for 2 days and their body weight and behavior were closely monitored during the 4 days following the procedure. Then, they were housed in groups of 2 mice per cage and behavioral tests started 10 days after surgery, enabling optimal recovery. 0.3μl of T3 (3,3’,5-Triiodo-L-thyronin, Sigma-T2877) or its vehicle was injected bilaterally in the dorsal CA1 (4ng/µl per side dissolved in 0.9% saline solution), 15min before starting the ORM training session, using silicone tubing connected to a peristaltic pump (PHD22/2000 Syringe Pump Infusion, Harvard Apparatus, Massachusetts, USA, flow rate: 0.1μl/min) and protruding 1 mm below cannulae tip. After behavior, the placement of cannulae in dorsal CA1 region of the hippocampus was verified histologically.

### Tissue collection

Hippocampi collection for pS6TRAP-RNA-Seq were done in two batches. First, mice were euthanized by decapitation between ZT2 and ZT4 (light phase), either 1h after ORM training (ORM+1h group, n=6 per diet) or control mice from their home cage (home cage ZT3 group, n=6 per diet). Second, mice were euthanized by decapitation between ZT13 and ZT16 (dark phase) either 12h after ORM training (ORM+12h group, n=6 per diet) or control mice from their home cage (home cage ZT15 group, n=5-6 per diet). For each mouse, hippocampus was dissected out on ice (Helbling 2021)[57]. In LoBind tube, tissue was then snap-frozen by submersion in liquid nitrogen prior to storage in a -80°C freezer.

### Translating Ribosome Affinity Purification (TRAP) mRNA isolation

For pS6 RNA-sequencing (pS6-TRAP-Seq) experiments, hippocampi were removed from -80°C freezer on ice to prevent any thawing and immediately complemented with 1ml of cold homogenization buffer (10mM HEPES [pH7.4], 150mM KCl, 10mM MgCl_2_, 2mM DTT, 0.1 complete Protease Inhibitor Cocktail (Sigma, 11836170001), 100U/ml RNasin Ribonuclease Inhibitors (Promega, N2515), 100µg/ml Cycloheximide). Tissues were disrupted with TissueLyser (Qiagen) for 15s – 30Hz. Homogenates were transferred in a new 1.5ml tubes and centrifuged for 10 min at 1000 g at 4°C. The supernatants were transferred to a new tube and NP40 10% (90µl) was added and incubated on ice for 5 min with gentle mixing. Samples were centrifuged for 10min at 10000g at 4°C. 350µl of RLT buffer (Qiagen, 74034) was added to 50µl of supernatants (as input samples), homogenates vortexed for 20-30s and snap-frozen in liquid nitrogen prior to storage in -80°C freezer until RNA purification. RNA from Input samples, reflecting hippocampus whole transcriptome, were purified and used to confirm pS6-TRAP enrichment by qPCR (data not shown). 500µl of supernatant was incubated with 1:25 anti-pS6 244-247 (ThermoFisher, 44-923G) (20µl) and incubated for 1,5 hour at 4°C with constant rotation. During anti-pS6 incubation, affinity purification beads were prepared where 200µl of Protein G Dynabeads (Thermofisher, 10009D) were washed 3 times for 10min in 200µl of Beads Washing Buffer (10mM HEPES [pH7.4], 300mM KCl, 10mM MgCl_2_, 1% NP40) and finally incubated for 10min in 200µl of Supplemented Homogenization Buffer (homogenization buffer supplemented with 1% NP40). Using magnet, beads were collected and homogenate/anti-pS6 solutions were added to beads for 1h incubation at 4°C with constant rotation. Following incubation, RNA bound to beads were washed 4 times for 10min at 4°C in 200µl of Wash Buffer (10mM HEPES [pH7.4], 350mM KCl, 10mM MgCl_2_, 2% NP40, 2mM DTT, 100U/ml RNasin Ribonuclease Inhibitors, 100µg/ml Cycloheximide), during the final wash, beads were placed onto the magnet and moved to room temperature. After the final wash 350µl of RLT buffer was immediately added to beads. Sample tubes were removed from the magnet and vortexed vigorously for 30s and incubated for 10min at room temperature. After removing beads using the magnet, mRNA was purified using RNeasy PLUS Micro kit (Qiagen, 74034). RNA integrity was assessed with 1µl of each mRNA-purified sample using RNA 6000 Pico kit (Agilent, 5067-1513) on an Agilent Bionalyzer with RNA Integrity Number (RIN) score up to 9 and mRNA quantity was measured using Nanodrop spectrophotometer. An average of 150 ng of RNA per sample was obtained.

### RNA sequencing and data analysis

Pooled equimolar library was synthesized at the Transcriptome facility using 50ng of TRAP-mRNA using Illumina Stranded mRNA Prep (Illumina, 20040534) and IDT for Illumina RNA UD Indexes Set B and Set C (Illumina, 20040554 and 20040555). Profiling libraries were assessed with LabChip GX Touch HT Nucleic Acid Analyzer (PerkinElmer, Revvity), with HT DNA NGS 3K Reagent Kit (PerkinElmer, CLS960013) onto DNA X-Mark Chip (PerkinElmer, CLS144006). Then libraries were quantified by qPCR onto LightCycler 480 (Roche) with NEB Next Library Quant Kit for Illumina (NEB, E7630L) and KAPA Library Quantification Kit (Roche, KK4854). Finally, libraries were sequenced at the PGTB facility (doi:10.15454/1.5572396583599417E12) with NextSeq 2000 P3 reagents (100 cycles) (Illumina, 20040559) onto NextSeq 2000 sequencer platform. Paired-end sequencing reads of 50bp resultd in an average of 60 million paired-end reads. Raw reads were processed using trimgalore (v0.6.7), cutadapt (v3.4), mapped against *Mus musculus* GRCm39 using HISAT2 aligner (v2.2.0) and quality checked with MultiQC (v3.9.5) through workflow Nextflow (v21.10.6) and nf-core/rnaseq (v3.6) pipeline onto genotoul bioinformatics platform Toulouse (Bioinfo Genotoul, https://doi.org/10.15454/1.5572369328961167E12). For differential analysis, reads mapped to unique transcripts were counted with featureCounts (Subread v2.0.3) annotation module. With Rstudio (Rversion 4.2.2), raw counts were filtered to keep genes with at least 10 counts across 73 samples. Differential expressed genes (DEG) were determined by the R Bioconductor package DESeq2 (v1.38.1) [Love, M.I. 2014][58] using negative binomial distribution with a model grouping all 73 samples. For Weighted Correlation Network Analysis (WGCNA, v1.72-1) [Langfelder P, 2008][31], ORM+1h (n=6) and HC-ZT3 (n=6) groups raw counts for each diet were analyzed separately. For each diet, filtered reads were normalized using DEseq2 variance stabilizing transformation (vst) function and were filtered to keep only genes with a variance larger than 0.025. This resulted in 2147 genes, 2391 genes and 1476 genes respectively for NC ad lib, HFS ad lib and HFS TRF which were used to create modules-associated genes networks. Within each module per diet, module membership and the associated *p-*value was calculated. For differential exon usage (DEU) analysis, reads mapped to unique exon were counted with featureCounts, DEXseq R Bioconductor package (v1.44.0) [Anders S, 2012][59] was used to measure DEU as a surrogate to infer differential splicing events in RNA-Seq data. Pipeline analysis is detailed as supplemental information.

## STATISTICAL ANALYSIS

### For metabolic data, behavioral analyses and qPCR data

Data were analyzed in GraphPad Prism 10 (GraphPad Software, Inc., San Diego, CA) unless indicated otherwise. Figure error bars represent the standard error of the mean (SEM). For comparisons, t-tests were used to determine significant differences between means of two group values (eg NC ad lib vs. HFS ad lib) and when more than 2 groups by 2-way ANOVA with Diet and ORM as factors, (or diet and TRF) followed by Tukey post-hoc tests for multiple comparisons, or 3 way ANOVA when 3 factors such as diet, TRF and ORM. For the kinetics of body weight across time a mixed model was used under 3-way ANOVA because of missing values in the table; repeated values were used for weeks of diet exposure. To analyze daily patterns of energy metabolism and feeding, data were fitted to cosinor regressions (SigmaPlot v13, Systat software Inc., San Jose, CA, USA) as follows: [y=A+(B·cos(2·π· (x−C)/24))] where A is the mean level, B the amplitude, C the acrophase of the 24-h rhythm. Significance is indicated as follows: * = p<0.05, ** = p<0.01, *** = p<0.001, **** = p<0.0001.

### For RNA-seq data analysis

Default parameters were used for all bioinformatic tools unless specified. Differential gene expression from RNA-Seq data was determined by DEseq2 using the negative binomial distribution (Wald test) and genes were considered as significantly differentially expressed when the Benjamini-Hochberg adjusted *p*-values < 0.05 (=FDR: False Discovery rate). In DEXseq analysis, exons were considered significantly differentially used when adjusted *p*-values < 0.05. Functional enrichment analysis such as KEGG pathways and Gene Ontology terms were performed using the R Bioconductor package clusterProfiler (v4.6.2) using enrichKEGG and enrichGO functions. The function compareCluster was used to compare enriched functional categories of gene module. Relaxed significance threshold of adjusted *p*-value < 0.2 was used for discovery and pathway analyses. Cell-type enrichment analyses to get specific gene signatures in different brain cell types (neurons, endothelial cell, astrocytes, microglia, oligodendrocyte precursor cells, newly formed oligodendrocytes, and myelinating oligodendrocytes) was performed using RNA-sequencing database [Zhang Y, 2014][32] and Fisher exact test was used to test enrichment of cell type signatures in genes whin modules provided by WGCNA analysis in each diet. Resulting *p*-values were corrected for multiple tests using the Benjamini-Hochberg method and the significant threshold was set to p<0.05 [Pandey RS, 2023][33]. Pipeline analysis is detailed as supplemental information.

Supplementary Tables are available on Zenodo repository with the doi: 10.5281/zenodo.11077283

## Acknowledgements

The authors thank the animal facility CIRCE, especially Gregory Artaxet and Eva Bruchet from Nutrineuro lab. cDNA libraries were prepared at the Transcriptome facility with the help of Frederic Martins (University of Bordeaux, INSERM, PUMA, Neurocentre Magendie, Bordeaux, France). Sequencing was done with the help of Zoé Delporte (Univ. Bordeaux, INRAE, BIOGECO, Cestas, France).

## Funding

This work was supported by INRAE (to MPM & G.F.), Fondation de Recherche sur le Cerveau (GF & MPM) and French National Research Agency (ANR-21-CE14-0086 MAORI) to MPM.

## Disclosures

Authors declare no competing interests.

## Author contributions

Conceptualization: MPM, GF, EC, FJ; Methodology: JCH, EGD, EM, AF, EC, GF; Investigations: JCH, RG, PM, DC, IMB, AC, EC, MRG, MPM; Funding acquisition: MPM, GF; Project administration and supervision: MPM; Writing–original draft: MPM; Editing: GF, EC, FJ, EM, RG, MRG, AF, LC.

---

**Fig. S1:**
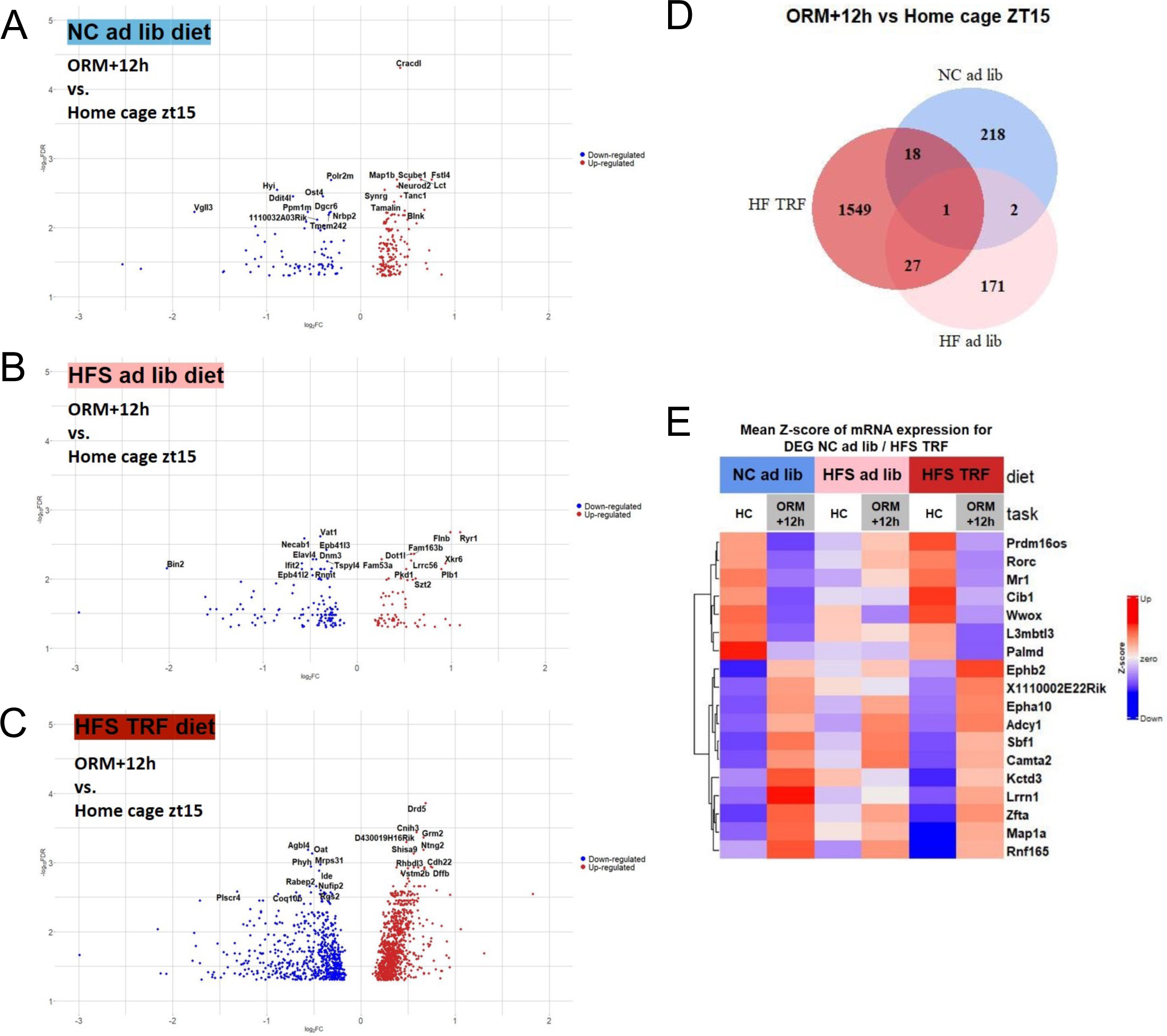
HFS diet moderately alters hippocampal translatome 12 hours after ORM training. A: Volcano plots and B: Venn diagram of DEG comparing ORM+12 vs zt15 groups. C: Heatmap of gene expression from DEG in common between NC and HFS TRF groups.

**Fig. S2:**
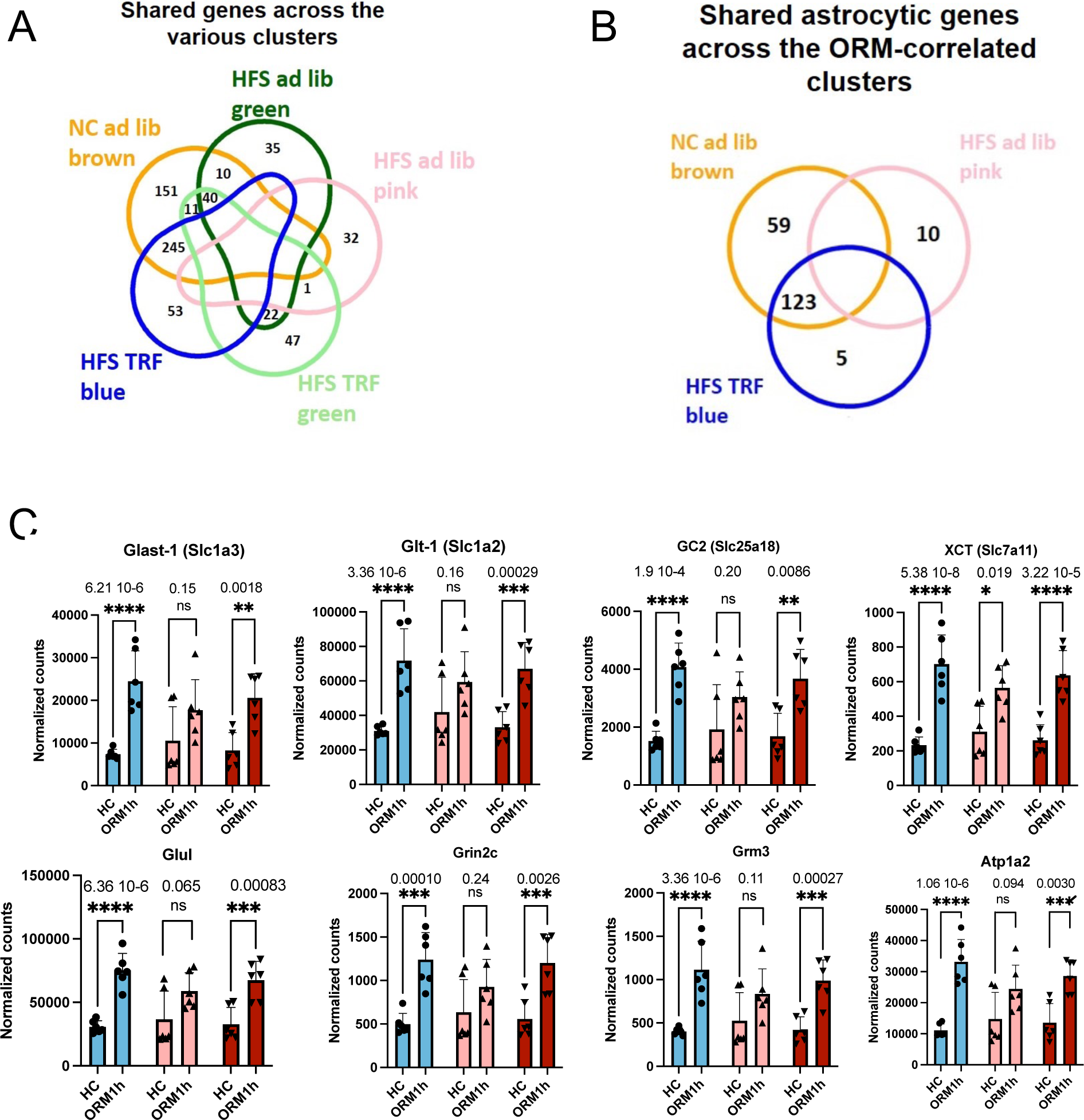
A: Venn diagram showing the number shared genes between the modules defined by the WGCNA analysis of Fig.4. The 245 genes in common between brown and blue module of respectively NC ad lib and HFS TRF were further examined in the study. B: Venn diagram showing the number of shared genes that are expressed in astrocytes and correlated with ORM in each diet. The 123 genes in common between brown and blue module of respectively NC ad lib and HFS TRF were further examined in the study.

**Fig. S3:**
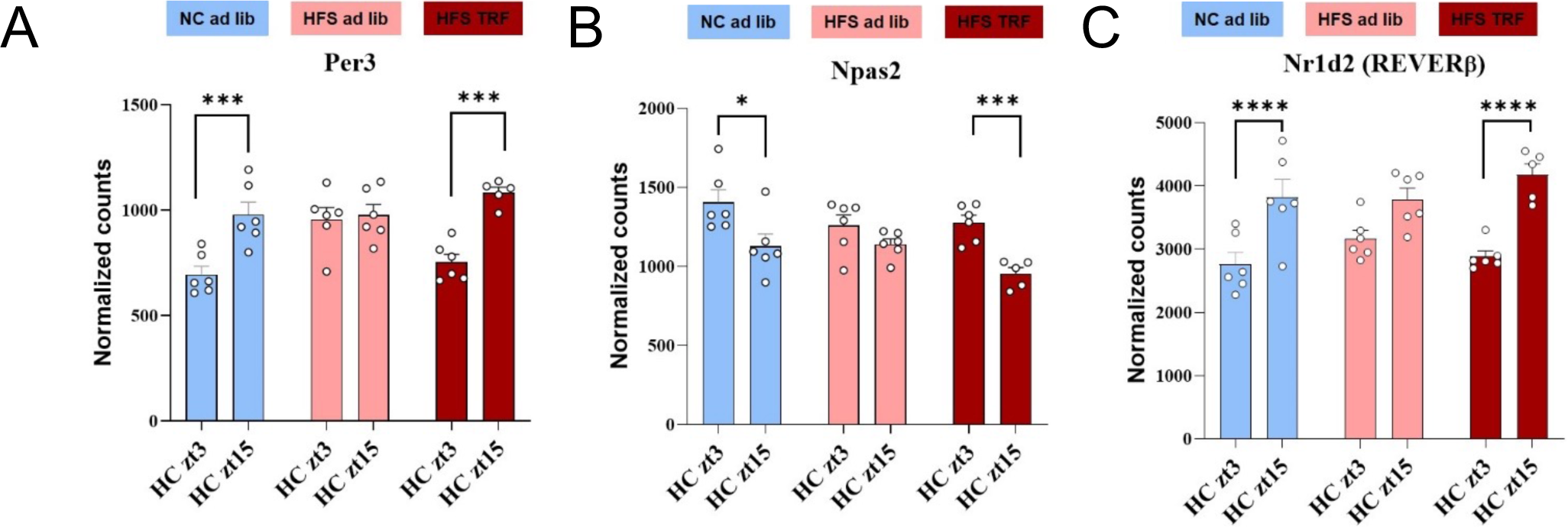
Gene expression of the core clock gene Per3, Npas2 and Nr1d2 from the HC zt15 vs HC zt3 comparison analyzed by DEseq2 package.

**Fig. S4:**
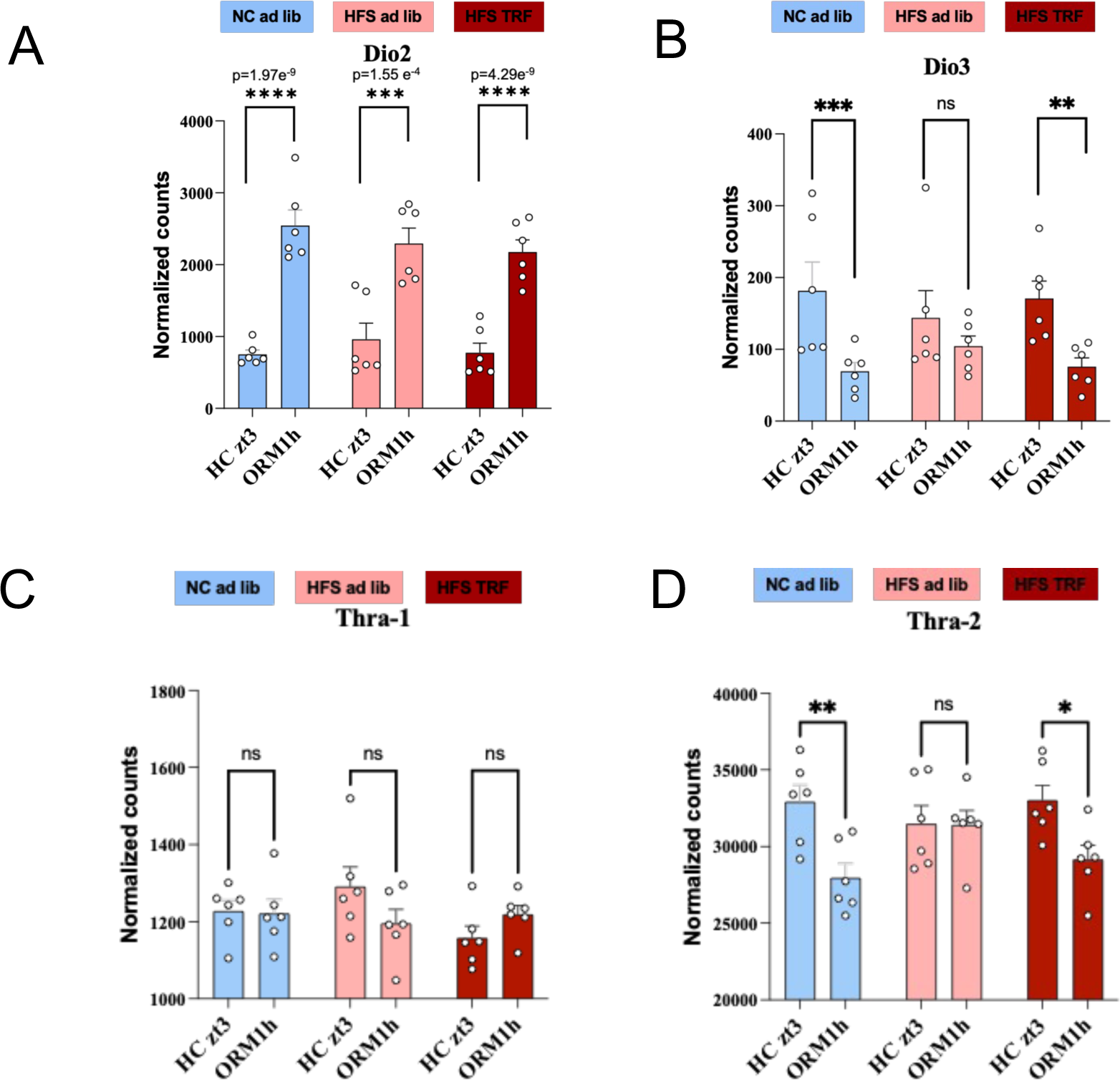
Gene expression of Dio2, Dio3, Thra1 and Thra2 from the ORM+1h vs. HC zt3 comparison analyzed by DEseq2 package.

## Notes

### Competing Interest Statement

The authors have declared no competing interest.

## References

[1] Lister NB, Baur LA, Felix JF, Hill AJ, Marcus C, Reinehr T, et al. Child and adolescent obesity. Nat Rev Dis Primers 2023;9:24. 10.1038/s41572-023-00435-4.

[2] Varela P, De Rosso S, Ferreira Moura A, Galler M, Philippe K, Pickard A, et al. Bringing down barriers to children’s healthy eating: a critical review of opportunities, within a complex food system. Nutr Res Rev 2023:1–21. 10.1017/S0954422423000203.

[3] Francis H, Stevenson R. The longer-term impacts of Western diet on human cognition and the brain. Appetite 2013;63:119–28. S0195-6663(12)00514-4 [pii] 10.1016/j.appet.2012.12.018.

[4] Sui SX, Pasco JA. Obesity and Brain Function: The Brain-Body Crosstalk. Medicina (Kaunas) 2020;56:499. 10.3390/medicina56100499.

[5] Del Olmo N, Ruiz-Gayo M. Influence of High-Fat Diets Consumed During the Juvenile Period on Hippocampal Morphology and Function. Front Cell Neurosci 2018;12:439. 10.3389/fncel.2018.00439.

[6] Morin J-P, Rodríguez-Durán LF, Guzmán-Ramos K, Perez-Cruz C, Ferreira G, Diaz-Cintra S, et al. Palatable Hyper-Caloric Foods Impact on Neuronal Plasticity. Frontiers in Behavioral Neuroscience 2017;11:19. 10.3389/fnbeh.2017.00019.

[7] Noble EE, Kanoski SE. Early life exposure to obesogenic diets and learning and memory dysfunction. Curr Opin Behav Sci 2016;9:7–14. 10.1016/j.cobeha.2015.11.014.

[8] Boitard C, Etchamendy N, Sauvant J, Aubert A, Tronel S, Marighetto A, et al. Juvenile, but not adult exposure to high-fat diet impairs relational memory and hippocampal neurogenesis in mice. Hippocampus 2012;22:2095–100. 10.1002/hipo.22032.

[9] Boitard C, Parkes SL, Cavaroc A, Tantot F, Castanon N, Layé S, et al. Switching Adolescent High-Fat Diet to Adult Control Diet Restores Neurocognitive Alterations. Frontiers in Behavioral Neuroscience 2016;10:225. 10.3389/fnbeh.2016.00225.

[10] Biyong EF, Alfos S, Dumetz F, Helbling J-C, Aubert A, Brossaud J, et al. Dietary vitamin A supplementation prevents early obesogenic diet-induced microbiota, neuronal and cognitive alterations. Int J Obes (Lond) 2021;45:588–98. 10.1038/s41366-020-00723-z.

[11] Naneix F, Bakoyiannis I, Santoyo-Zedillo M, Bosch-Bouju C, Pacheco-Lopez G, Coutureau E, et al. Chemogenetic silencing of hippocampus and amygdala reveals a double dissociation in periadolescent obesogenic diet-induced memory alterations. Neurobiol Learn Mem 2021;178:107354. 10.1016/j.nlm.2020.107354.

[12] Vouimba R-M, Bakoyiannis I, Ducourneau E-G, Maroun M, Ferreira G. Bidirectional modulation of hippocampal and amygdala synaptic plasticity by post-weaning obesogenic diet intake in male rats: Influence of the duration of diet exposure. Hippocampus 2021;31:117–21. 10.1002/hipo.23278.

[13] Bakoyiannis I, Ducourneau EG, N’diaye M, Fermigier A, Ducroix-Crepy C, Bosch-Bouju C, et al. Obesogenic diet induces circuit-specific memory deficits in mice. Elife 2024;13:e80388. 10.7554/eLife.80388.

[14] Kohsaka A, Laposky AD, Ramsey KM, Estrada C, Joshu C, Kobayashi Y, et al. High-fat diet disrupts behavioral and molecular circadian rhythms in mice. Cell Metabolism 2007;6:414–21. 10.1016/j.cmet.2007.09.006.

[15] Mendoza J, Pévet P, Challet E. High-fat feeding alters the clock synchronization to light. The Journal of Physiology 2008;586:5901–10. 10.1113/jphysiol.2008.159566.

[16] Challet E. Circadian clocks, food intake, and metabolism. Progress in Molecular Biology and Translational Science 2013;119:105–35. 10.1016/B978-0-12-396971-2.00005-1.

[17] Asher G, Sassone-Corsi P. Time for food: the intimate interplay between nutrition, metabolism, and the circadian clock. Cell 2015;161:84–92. S0092-8674(15)00302-5 [pii] 10.1016/j.cell.2015.03.015.

[18] Eckel-Mahan KL, Patel VR, de Mateo S, Orozco-Solis R, Ceglia NJ, Sahar S, et al. Reprogramming of the circadian clock by nutritional challenge. Cell 2013;155:1464–78. S0092-8674(13)01485-2 [pii] 10.1016/j.cell.2013.11.034.

[19] Tognini P, Samad M, Kinouchi K, Liu Y, Helbling J-C, Moisan M-P, et al. Reshaping circadian metabolism in the suprachiasmatic nucleus and prefrontal cortex by nutritional challenge. Proc Natl Acad Sci U S A 2020;117:29904–13. 10.1073/pnas.2016589117.

[20] Boege HL, Bhatti MZ, St-Onge M-P. Circadian rhythms and meal timing: impact on energy balance and body weight. Curr Opin Biotechnol 2020;70:1–6. 10.1016/j.copbio.2020.08.009.

[21] Manoogian ENC, Chow LS, Taub PR, Laferrère B, Panda S. Time-restricted Eating for the Prevention and Management of Metabolic Diseases. Endocr Rev 2022;43:405–36. 10.1210/endrev/bnab027.

[22] Stucchi P, Gil-Ortega M, Merino B, Guzman-Ruiz R, Cano V, Valladolid-Acebes I, et al. Circadian feeding drive of metabolic activity in adipose tissue and not hyperphagia triggers overweight in mice: is there a role of the pentose-phosphate pathway? Endocrinology 2012;153:690–9. en.2011-1023 [pii] 10.1210/en.2011-1023.

[23] Hatori M, Vollmers C, Zarrinpar A, DiTacchio L, Bushong EA, Gill S, et al. Time-restricted feeding without reducing caloric intake prevents metabolic diseases in mice fed a high-fat diet. Cell Metab 2012;15:848–60. S1550-4131(12)00189-1 [pii] 10.1016/j.cmet.2012.04.019.

[24] Chaix A, Zarrinpar A, Miu P, Panda S. Time-restricted feeding is a preventative and therapeutic intervention against diverse nutritional challenges. Cell Metab 2014;20:991–1005. 10.1016/j.cmet.2014.11.001.

[25] Whittaker DS, Akhmetova L, Carlin D, Romero H, Welsh DK, Colwell CS, et al. Circadian modulation by time-restricted feeding rescues brain pathology and improves memory in mouse models of Alzheimer’s disease. Cell Metab 2023;35:1704–1721.e6. 10.1016/j.cmet.2023.07.014.

[26] Davis JA, Paul JR, Yates SD, Cutts EJ, McMahon LL, Pollock JS, et al. Time-restricted feeding rescues high-fat-diet-induced hippocampal impairment. iScience 2021;24:102532. 10.1016/j.isci.2021.102532.

[27] Delorme J, Wang L, Kodoth V, Wang Y, Ma J, Jiang S, et al. Hippocampal neurons’ cytosolic and membrane-bound ribosomal transcript profiles are differentially regulated by learning and subsequent sleep. Proc Natl Acad Sci U S A 2021;118:e2108534118. 10.1073/pnas.2108534118.

[28] Delorme J, Wang L, Kuhn FR, Kodoth V, Ma J, Martinez JD, et al. Sleep loss drives acetylcholine-and somatostatin interneuron-mediated gating of hippocampal activity to inhibit memory consolidation. Proc Natl Acad Sci U S A 2021;118:e2019318118. 10.1073/pnas.2019318118.

[29] Baghcheghi Y, Salmani H, Beheshti F, Hosseini M. Contribution of Brain Tissue Oxidative Damage in Hypothyroidism-associated Learning and Memory Impairments. Adv Biomed Res 2017;6:59. 10.4103/2277-9175.206699.

[30] Rial-Pensado E, Canaple L, Guyot R, Clemmensen C, Wiersema J, Wu S, et al. Neuronal Blockade of Thyroid Hormone Signaling Increases Sensitivity to Diet-Induced Obesity in Adult Male Mice. Endocrinology 2023;164:bqad034. 10.1210/endocr/bqad034.

[31] Langfelder P, Horvath S. WGCNA: an R package for weighted correlation network analysis. BMC Bioinformatics 2008;9:559. 10.1186/1471-2105-9-559.

[32] Zhang Y, Chen K, Sloan SA, Bennett ML, Scholze AR, O’Keeffe S, et al. An RNA-sequencing transcriptome and splicing database of glia, neurons, and vascular cells of the cerebral cortex. J Neurosci 2014;34:11929–47. 10.1523/JNEUROSCI.1860-14.2014.

[33] Pandey RS, Kotredes KP, Sasner M, Howell GR, Carter GW. Differential splicing of neuronal genes in a Trem2*R47H mouse model mimics alterations associated with Alzheimer’s disease. BMC Genomics 2023;24:172. 10.1186/s12864-023-09280-x.

[34] Zekri Y, Guyot R, Flamant F. An Atlas of Thyroid Hormone Receptors’ Target Genes in Mouse Tissues. Int J Mol Sci 2022;23:11444. 10.3390/ijms231911444.

[35] Whittaker DS, Loh DH, Wang H-B, Tahara Y, Kuljis D, Cutler T, et al. Circadian-based Treatment Strategy Effective in the BACHD Mouse Model of Huntington’s Disease. J Biol Rhythms 2018;33:535–54. 10.1177/0748730418790401.

[36] Hepler C, Weidemann BJ, Waldeck NJ, Marcheva B, Cedernaes J, Thorne AK, et al. Time-restricted feeding mitigates obesity through adipocyte thermogenesis. Science 2022;378:276–84. 10.1126/science.abl8007.

[37] Pendergast JS, Branecky KL, Yang W, Ellacott KLJ, Niswender KD, Yamazaki S. High-fat diet acutely affects circadian organisation and eating behavior. The European Journal of Neuroscience 2013;37:1350–6. 10.1111/ejn.12133.

[38] Gallop MR, Tobin SY, Chaix A. Finding balance: understanding the energetics of time-restricted feeding in mice. Obesity (Silver Spring) 2023;31 Suppl 1:22–39. 10.1002/oby.23607.

[39] Alberini CM, Kandel ER. The regulation of transcription in memory consolidation. Cold Spring Harbor Perspectives in Biology 2014;7:a021741. 10.1101/cshperspect.a021741.

[40] Oliveira AMM. DNA methylation: a permissive mark in memory formation and maintenance. Learning & Memory (Cold Spring Harbor, NY) 2016;23:587–93. 10.1101/lm.042739.116.

[41] Kwapis JL, Alaghband Y, Kramár EA, López AJ, Vogel Ciernia A, White AO, et al. Epigenetic regulation of the circadian gene Per1 contributes to age-related changes in hippocampal memory. Nature Communications 2018;9:3323. 10.1038/s41467-018-05868-0.

[42] Sui L, Wang F, Liu F, Wang J, Li BM. Dorsal hippocampal administration of triiodothyronine enhances long-term memory for trace cued and delay contextual fear conditioning in rats. J Neuroendocrinol 2006;18:811–9. 10.1111/j.1365-2826.2006.01480.x.

[43] Sui L, Ren W-W, Li B-M. Administration of thyroid hormone increases reelin and brain-derived neurotrophic factor expression in rat hippocampus in vivo. Brain Res 2010;1313:9–24. 10.1016/j.brainres.2009.12.010.

[44] Perea G, Navarrete M, Araque A. Tripartite synapses: astrocytes process and control synaptic information. Trends Neurosci 2009;32:421–31. 10.1016/j.tins.2009.05.001.

[45] Valtcheva S, Venance L. Control of Long-Term Plasticity by Glutamate Transporters. Front Synaptic Neurosci 2019;11:10. 10.3389/fnsyn.2019.00010.

[46] Tsai S-F, Hsu P-L, Chen Y-W, Hossain MS, Chen P-C, Tzeng S-F, et al. High-fat diet induces depression-like phenotype via astrocyte-mediated hyperactivation of ventral hippocampal glutamatergic afferents to the nucleus accumbens. Mol Psychiatry 2022;27:4372–84. 10.1038/s41380-022-01787-1.

[47] Lau BK, Murphy-Royal C, Kaur M, Qiao M, Bains JS, Gordon GR, et al. Obesity-induced astrocyte dysfunction impairs heterosynaptic plasticity in the orbitofrontal cortex. Cell Rep 2021;36:109563. 10.1016/j.celrep.2021.109563.

[48] Valladolid-Acebes I, Merino B, Principato A, Fole A, Barbas C, Lorenzo MP, et al. High-fat diets induce changes in hippocampal glutamate metabolism and neurotransmission. Am J Physiol Endocrinol Metab 2012;302:E396–402. 10.1152/ajpendo.00343.2011.

[49] Karimi SA, Komaki A, Salehi I, Sarihi A, Shahidi S. Role of group II metabotropic glutamate receptors (mGluR2/3) blockade on long-term potentiation in the dentate gyrus region of hippocampus in rats fed with high-fat diet. Neurochem Res 2015;40:811–7. 10.1007/s11064-015-1531-3.

[50] Morte B, Manzano J, Scanlan TS, Vennström B, Bernal J. Aberrant maturation of astrocytes in thyroid hormone receptor alpha 1 knockout mice reveals an interplay between thyroid hormone receptor isoforms. Endocrinology 2004;145:1386–91. 10.1210/en.2003-1123.

[51] Mendes-de-Aguiar CBN, Alchini R, Decker H, Alvarez-Silva M, Tasca CI, Trentin AG. Thyroid hormone increases astrocytic glutamate uptake and protects astrocytes and neurons against glutamate toxicity. J Neurosci Res 2008;86:3117–25. 10.1002/jnr.21755.

[52] Paisdzior S, Knierim E, Kleinau G, Biebermann H, Krude H, Straussberg R, et al. A New Mechanism in THRA Resistance: The First Disease-Associated Variant Leading to an Increased Inhibitory Function of THRA2. Int J Mol Sci 2021;22:5338. 10.3390/ijms22105338.

[53] Gould E, Frankfurt M, Westlind-Danielsson A, McEwen BS. Developing forebrain astrocytes are sensitive to thyroid hormone. Glia 1990;3:283–92. 10.1002/glia.440030408.

[54] Vidmar AP, Naguib M, Raymond JK, Salvy SJ, Hegedus E, Wee CP, et al. Time-Limited Eating and Continuous Glucose Monitoring in Adolescents with Obesity: A Pilot Study. Nutrients 2021;13:3697. 10.3390/nu13113697.

[55] Naguib MN, Hegedus E, Raymond JK, Goran MI, Salvy S-J, Wee CP, et al. Continuous Glucose Monitoring in Adolescents With Obesity: Monitoring of Glucose Profiles, Glycemic Excursions, and Adherence to Time Restricted Eating Programs. Front Endocrinol (Lausanne) 2022;13:841838. 10.3389/fendo.2022.841838.

[56] Leger M, Quiedeville A, Bouet V, Haelewyn B, Boulouard M, Schumann-Bard P, et al. Object recognition test in mice. Nature Protocols 2013;8:2531–7. 10.1038/nprot.2013.155.

[57] Helbling JC, Kinouchi K, TrifilieffP, Sassone-Corsi P, Moisan MP. Combined Gene Expression and Chromatin Immunoprecipitation From a Single Mouse Hippocampus. Current Protocols 2021;1.

[58] Love MI, Huber W, Anders S. Moderated estimation of fold change and dispersion for RNA-seq data with DESeq2. Genome Biol 2014;15:550. 10.1186/s13059-014-0550-8.

[59] Anders S, Reyes A, Huber W. Detecting differential usage of exons from RNA-seq data. Genome Res 2012;22:2008–17. 10.1101/gr.133744.111.

